# Targeting riboswitches with beta-axial substituted cobalamins

**DOI:** 10.1101/2022.12.25.521693

**Authors:** Shelby R. Lennon, Aleksandra J. Wierzba, Shea H. Siwik, Dorota Gryko, Amy E. Palmer, Robert T. Batey

**Affiliations:** Department of Biochemistry, University of Colorado, Boulder, CO 80309-0596, USA; BioFrontiers Institute, University of Colorado, Boulder, CO 80303 – 0596, USA; Institute of Organic Chemistry, Polish Academy of Sciences, Kasprzaka 44/52, 01-224 Warsaw, Poland

## Abstract

RNA-targeting small molecule therapeutics an emerging field hindered by an incomplete understanding of the basic principles governing RNA-ligand interactions. One way to advance our knowledge in this area is to study model systems where these interactions are better understood, such as riboswitches. Riboswitches bind a wide array of small molecules with high affinity and selectivity, providing a wealth of information on how RNA recognizes ligands through diverse structures. The cobalamin-sensing riboswitch is a particularly useful model system as similar sequences show highly specialized binding preferences for different biological forms of cobalamin. This riboswitch is also widely dispersed across bacteria and therefore holds strong potential as an antibiotic target. Many synthetic cobalamin forms have been developed for various purposes including therapeutics, but their interaction with cobalamin riboswitches is yet to be explored. In this study, we characterize the interactions of eleven cobalamin derivatives with three representative cobalamin riboswitches using *in vitro* binding experiments (both chemical footprinting and a fluorescence-based assay) and a cell-based reporter assay. The derivatives show productive interactions with two of the three riboswitches, demonstrating simultaneously plasticity and selectivity within these RNAs. The observed plasticity is partially achieved through a novel structural rearrangement within the ligand binding pocket, providing insight into how similar RNA structures can be targeted in the future. As the derivatives also show *in vivo* functionality, they serve as several potential lead compounds for further drug development.

## Introduction

RNA presents an underutilized target for therapeutics with the potential for wide-reaching effects.^3-5^ While drugs to combat disease have historically targeted proteins, the revelation that only about 1.5% of the human genome is translated into protein^6^ while at least 75% is transcribed into RNA^7^ suggests that an RNA-focused approach may provide a wealth of targeting opportunities.^8-9^ However, beyond antibacterial and antiviral drugs^10-11^, small molecule-based RNA targeting therapeutics are mostly in the early discovery stages of development.^3^ One of the main challenges is that our understanding of the basic principles of RNA-small molecule interactions is lacking as compared to that of proteins.^4-5, 12^ While many nonspecific RNA binding small molecules, such as general intercalators, are known, effective therapeutics need to be specific for a target RNA sequence or structure. Further exploration of t how RNA specifically recognizes and binds various ligands will facilitate rational design of small molecules to target and drug RNA.

Perhaps no system better demonstrates specific and high affinity RNA-small molecule interactions that drive a biological outcome than riboswitches. Riboswitches are non-coding RNA elements typically found in the 5’-leader of messenger RNAs (mRNAs), mostly in bacteria.^13^ Small molecule binding to the riboswitch regulates expression of the mRNA by directing formation of alternative RNA structures that inform the expression machinery. To date, over 55 classes of riboswitches have been discovered, binding a wide range of metabolites.^13^ These ligands can be related to RNA structurally or metabolically, such as nucleotide derivatives, but can also be non-self-compounds, such as elemental ions and amino acids such as lysine. Riboswitches can adopt a wide spectrum of RNA structures to create binding pockets with high affinity and specificity for these target ligands. These structures range from simple sites comprising formation of base triples with the ligand^14-15^ to complex multi-junction pockets.^2, 16^ Riboswitch binding pockets often contain common motifs such as pseudoknots, suggesting that similar motifs in other RNAs can also be targeted by drug-like small molecules. Thus, the diversity of small molecules and RNA structures encompassed by riboswitches presents a rich palette of robust model systems for the study of RNA-small molecule interactions, and several studies have shown the strength of using riboswitches in this manner.^17-20^

The cobalamin (Cbl, B_12_) riboswitch controls the biosynthesis, salvage, and transport of Cbl, an essential metabolite for many forms of life.^21^ The genes which maintain Cbl cellular levels can be essential for survival and virulence of bacterial pathogens, making the Cbl riboswitch a promising antibiotic target.^22^ Corresponding to the importance of this metabolite, the Cbl riboswitch is the second most widely distributed, found in almost every major clade of bacteria.^23^ The Cbl riboswitch is a particularly useful model system for the study of RNA-small molecule interactions, as different classes show highly specialized binding to the biological forms of Cbl despite similar sequences. Currently two classes, I and II, are recognized within the Cbl riboswitch family. Class-I specifically binds 5’-deoxyadenosyl-Cbl (AdoCbl) and class-II exhibits a spectrum of selectivity for AdoCbl and methylcobalamin (MeCbl).^24^ All forms of Cbl contain a corrin ring that coordinates a cobalt atom and is flanked by methyl, acetamide, and propionamide groups. The sixth coordination site of the cobalt ion is the beta-axial position (**Figure 1A**), coordinating either the 5’-deoxyadenosyl (Ado) or methyl group. The interaction of the beta-axial group with a universally conserved four-way junction determines the selectivity of the riboswitch.^24^ Understanding how similar RNA structures can bind distinct compounds and how that can be exploited using analogs can advance the optimization of lead compounds in drug discovery.

**Figure 1.**
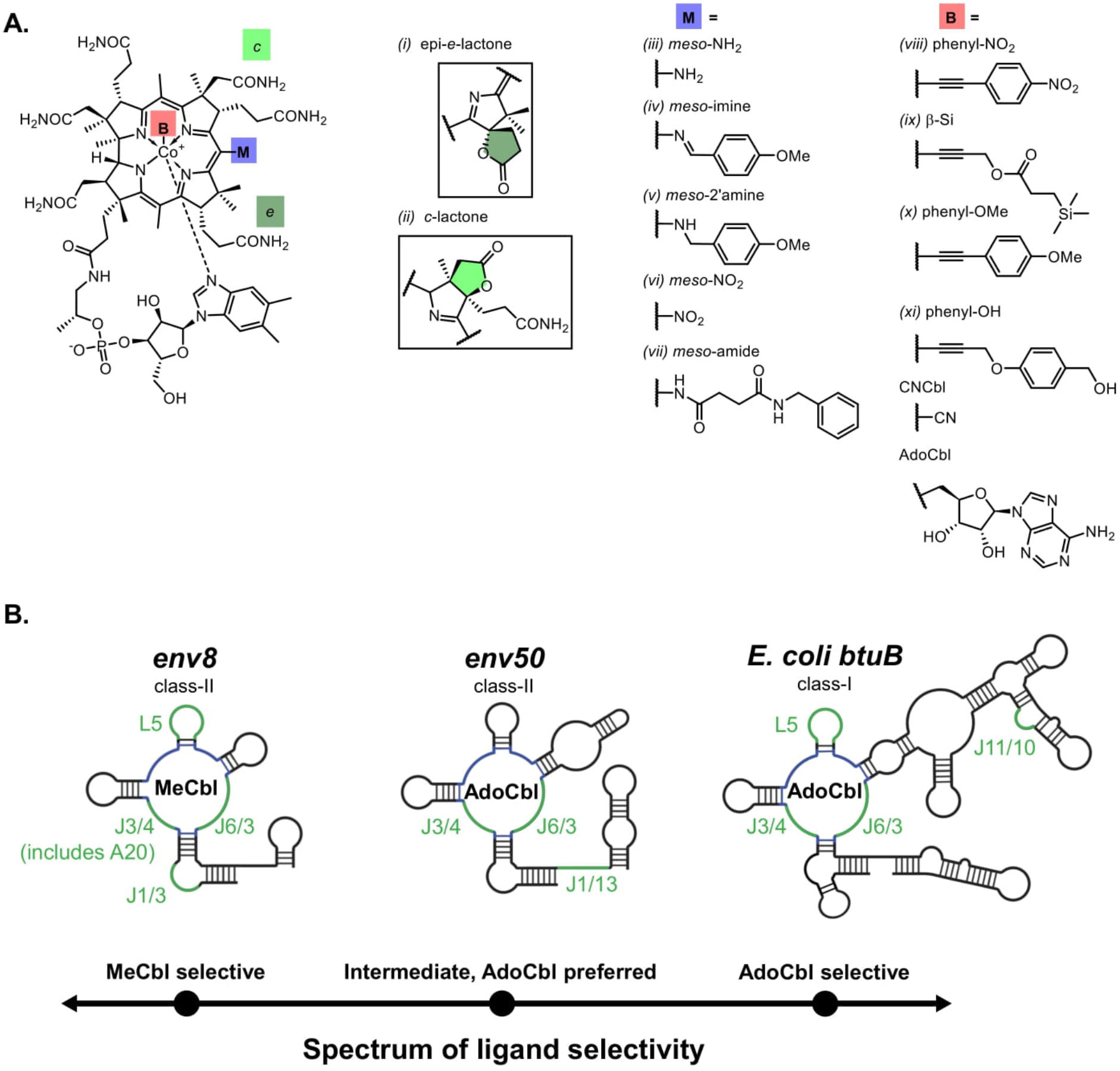
Structures of riboswitches and Cbl derivatives used in this study. (**A**) Chemical structures of the Cbl derivatives, identified by Roman numerals. The modifications for the lactone derivatives are shown as insets, with the lactone groups at the *e* and *c* positions. For the *meso* and beta-axial derivatives, only the modifications are shown, with the positions marked on the Cbl as “M” and “B” respectively. There is a cyano group at the beta-axial position for the *meso* and lactone derivatives. There is a hydrogen at the *meso* position for the beta axial and lactone derivatives. Also included are the beta-axial groups for AdoCbl and CNCbl. (**B**) Secondary structures of the Cbl riboswitches. The preferred binding forms of Cbl are written in the common central four-way junction of each structure, colored in blue. Colored and labeled in green are additional areas relevant to this paper. Created using Biorender.com.

Synthetic forms of Cbl have also been developed for various purposes, including therapeutics.^25^ Cyanocobalamin (CNCbl) has a light-stable cyano group in the beta-axial position (**Figure 1A**) and is used as a diet supplement as it can be converted to the natural forms within the body. Cbl derivatives that cannot be transformed into the biologically active forms have also been developed and are called antivitamins.^25^ Like CNCbl, B_12_ antivitamins can have modifications at the beta-axial position. Additional modifications include formation of a lactone group at the corrin ring *c* position.^26^ B_12_ antivitamins are often used to study B_12_ deficiency and have potential as novel antibiotics or cancer drugs.^27-28^ *Meso*-modified Cbl derivatives have also been developed as delivery systems for drugs, imaging agents, oligonucleotides, and other agents.^29-31^ These derivatives have modified chemical properties, caused by addition of electron-donating or -withdrawing groups around the corrin ring to affect the redox potential of the central cobalt ion.^32^ The interaction of these synthetic Cbl forms with Cbl binding proteins has been studied.^33-34^ However, the potential use of structurally modified cobalamins as antibiotics through interaction with Cbl riboswitches has only been hypothesized^25, 27^ as their ability to bind to Cbl riboswitches has not been studied.

Here we directly address this question by examining the interactions of a series of Cbl derivatives with representative Cbl riboswitches. Three Cbl riboswitches were chosen to represent the structural and functional diversity across this family of RNAs:^35^ the class-I *E. coli btuB* riboswitch that specifically binds AdoCbl, a class-II riboswitch that has high selectivity for small beta-axial forms of Cbl (*env8*), and a class-II riboswitch that has moderate selectivity for large beta-axial form Cbl but also binds small beta-axial cobalamins (*env50*) (**Figure 1B**). Eleven Cbl derivatives (**Figure 1A**) synthesized with many of the modifications discussed above were compared to AdoCbl and CNCbl in binding assays. These modifications include the presence of lactone groups at the *e* and *c* corrin ring positions (derivatives *i* and *ii)*, various modifications to the *meso* position (derivatives *iii* through *vii*), and substitutions at the beta-axial position (derivatives *viii* through *xi*). Each ligand’s binding properties were evaluated using chemical probing and a complementary fluorescent displacement assay. The results reveal surprising plasticity of the class-II riboswitches, including a novel binding mode to accommodate bulky beta-axial moieties. The interactions are shown to be biologically relevant using an *in vivo* cell reporter assay, establishing some of the derivatives as promising lead compounds for drug development.

## Results and Discussion

### Bulky beta-axial substitutions displace a purine residue in the binding pocket of a MeCbl-selective riboswitch

To assess the potential of the above Cbl derivative panel to bind Cbl riboswitches, we docked a subset onto the *env8* crystal structure (PDB ID 4FRN).^2^ If the RNA surrounding the binding pocket is rigid and the derivatives interact similarly to CNCbl, then this analysis suggests that these derivatives would be incapable of binding. Each modification to the corrin ring appears to sterically clash with key RNA architectural elements as the corrin ring and beta-axial face are buried within the core of the riboswitch. The beta-axial derivatives in particular present an obstacle with their bulky moieties due to clashes with nucleotides A20 (J3/4) and A68 (J6/3) in the most conserved region of the binding pocket (**Figure S1**). The lactone and *meso* derivatives clash with nucleotides G70 and C71 (J6/3). However, studies with other riboswitches have revealed unexpected modes of plasticity in the RNA that enable binding of chemical analogs.^16, 36-38^

To visualize binding of Cbl derivatives to the three representative riboswitches, we used selective 2’-hydroxyl acylation analyzed by primer extension (SHAPE)^39^, which has previously been used to characterize Cbl riboswitches.^2, 40^ In support of conformational plasticity in the cobalamin riboswitch, SHAPE revealed that many of the tested Cbl derivatives bind to *env8* (**Figure 2A,B and S2**). CNCbl was used as a standard for chemical reactivity protections and enhancements in the RNA in response to ligand binding. The presence of CNCbl promoted the previously observed set of reactivity changes in L5, J6/3, and J1/13 that are indicative of binding.^40^ The derivatives promoted the same protections and deprotections in the *env8* riboswitch to varying extents. Notably, the reactivity pattern of the beta-axial derivatives was similar to that of CNCbl, despite the predicted steric clash. The lactone and *meso* derivatives displayed a larger range of induced reactivity changes, with some behaving similarly to CNCbl and several similar to the no ligand condition, suggesting a range of binding affinities. The lack of significant differences between the footprinting patterns for CNCbl and most of the derivatives suggests that they all interact with the RNA in a similar fashion, indicating that the RNA can accommodate many of these modifications without large structural rearrangements.

**Figure 2.**
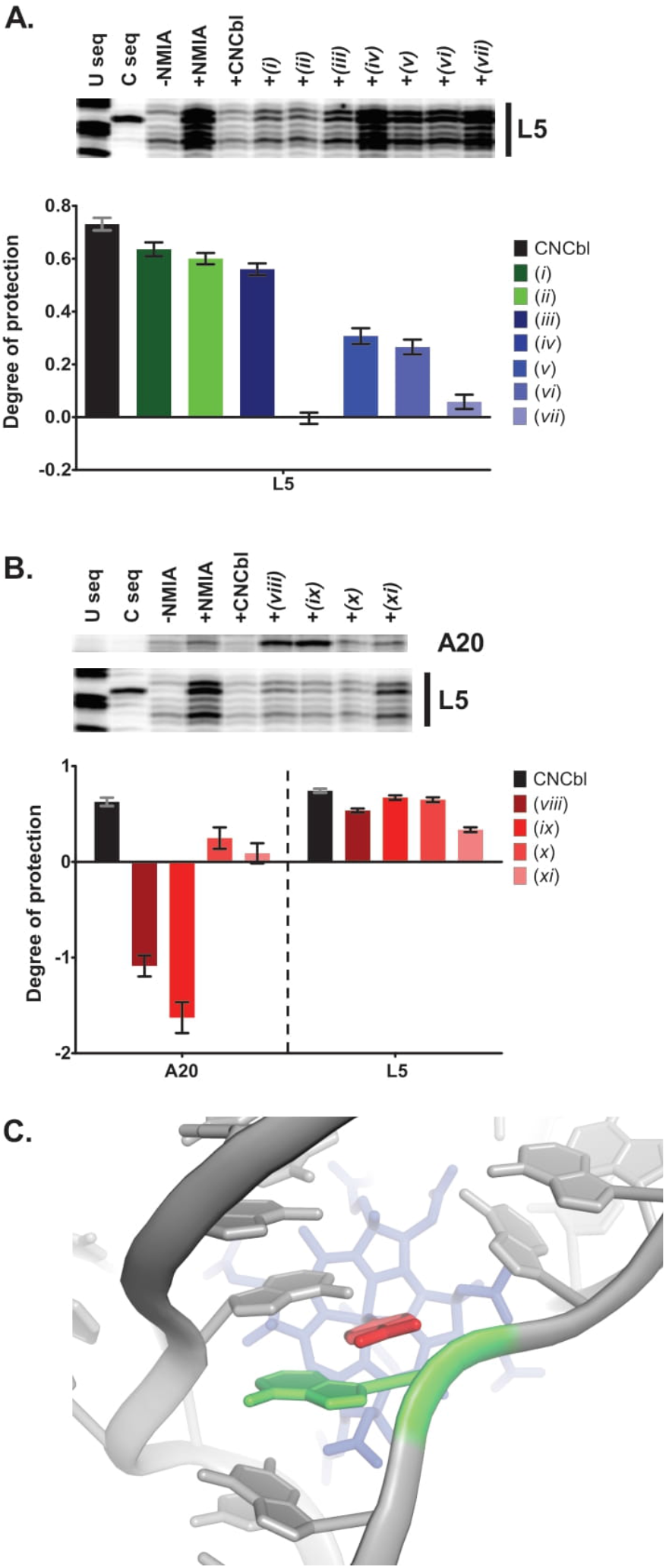
Structural probing by SHAPE of Cbl derivatives binding to *env8*. Representative gels are shown and the results are quantified (n=6-10, error is reported as SEM). A) Lactone and *meso* derivatives, highlighting the L5 region. B) Same as A, but with the beta-axial derivatives. Nucleotide A20 is also included. C) Crystal structure of *env8* (PDB ID 4FRG)^2^ with phenyl-NO_2_ Cbl derivative (*viii*, Cambridge Structural Database deposition number 961228), superimposed over CNCbl (not shown) to show steric clash with position A20. *env8* is colored gray, the base Cbl molecule blue, the phenyl-NO_2_ moiety red, and A20 green.

Only nucleotide A20 displayed a significant difference in chemical reactivity for several derivatives as compared to CNCbl. Whereas CNCbl binding protects this position, binding of beta-axial derivatives *viii* and *ix* show a strong enhancement of chemical reactivity. A20 is in a region with structural significance, being one of the four nucleotides from J3/4 and J6/3 (G19, A20, A68, and A67) that comprise a cross-strand purine stack^41-42^ forming one half of the ligand binding pocket (**Figure 2C**). The beta-axial face of CNCbl forms direct van der Waals contacts with this region of the RNA. We propose that the nucleotide deprotection is the result of A20 being displaced from the purine stack by the derivative beta-axial group. This is similar to the observed displacement of a key nucleotide in the purine riboswitch by guanine derivatives containing modifications at the N2 position.^36^ Removal of the nucleotide from the protective environment of the purine stack would enhance the reactivity of its 2’-hydroxyl group towards the probing reagent and enable the bulky beta-axial group to intercalate between residues G19 and A69.

Intercalation of a functional group into the purine spine of Cbl riboswitches has been observed for AdoCbl binding.^2, 43-44^ In all AdoCbl-riboswitch structures, nucleotides from J3/4 and J6/3 form a continuous base-stack as one half of the binding pocket, and an opening within the stack allows for insertion of the Ado moiety. The opening is created by J6/3 being pulled back and away from the binding pocket, supported through interactions with another region of the RNA. In the class-I structures, the adenosine equivalent to A70 in J6/3 of *env8* forms a *trans* A·A Watson-Crick-Hoogsteen pairing with the nucleobase of the Ado moiety.^2, 44^ This nucleotide is not reactive in SHAPE analysis due to its interactions with the Ado.^2^ This contrasts with a class-II structure where a J6/3 nucleotide is displaced from the purine stack but does not interact with the Ado.^43^ Due to its selectivity against AdoCbl, it was unexpected that *env8* could act similarly to the AdoCbl riboswitches by binding bulky beta-axial groups that displace a nucleotide from the purine stack. This is also the first evidence of displacement of a nucleotide in J3/4 as opposed to J6/3. Thus, both halves of the riboswitch central spine display plasticity that can be exploited by Cbl derivatives with bulky beta-axial substitutions.

### Chemical probing suggests the tested Cbl derivatives do not bind to AdoCbl-selective riboswitches

Given the above considerations, we predicted that AdoCbl-selective riboswitches could bind to a set of the Cbl derivatives panel. The class-I *btuB* Cbl riboswitch does not productively bind Cbl forms with small axial ligands^45^ and therefore would not be expected to bind the lactone or *meso* Cbl derivatives, which have a cyano beta-axial moiety. In contrast, the class-II *env50* Cbl riboswitch binds both large and small beta-axial Cbl forms and hence might bind a similar set of derivatives as *env8*. Since *env50* has a 23-fold preference for binding AdoCbl over MeCbl,^24^ the beta-axial derivatives may have a higher binding affinity for *env50* than the lactone and *meso* derivatives.

To experimentally validate these predictions, we assessed derivative binding to *env50* and *btuB* by SHAPE. *Env50* showed protections in J6/3 and deprotections in J1/13 in the presence of AdoCbl, indicative of binding (**Figure 3A and Figure S3**). Unexpectedly, neither CNCbl nor any of the tested Cbl derivatives induced detectable pattern changes in the RNA despite the known binding of MeCbl to this riboswitch. While CNCbl may have a slightly different binding affinity to *env50* than MeCbl, it was still expected to exhibit a SHAPE signature, especially at high ligand concentration (30 µM). In the case of the *btuB* riboswitch, we observed a set of protections in L5 and J11/10 in the presence of AdoCbl, consistent with previous studies (**Figure 3B and Figure S4**).^2^ However, similar to *env50*, chemical probing in the presence of CNCbl or any of the panel of Cbl derivatives did not yield reactivity pattern differences in the RNA. While this was expected for the lactone and *meso* derivatives, these results were unexpected for the beta-axial derivatives. Thus, the SHAPE probing suggested neither *env50* nor *btuB* bind the panel of Cbl derivatives. The failure of SHAPE to detect binding may be a result of chemical footprinting being unable to detect weakly binding compounds.

**Figure 3.**
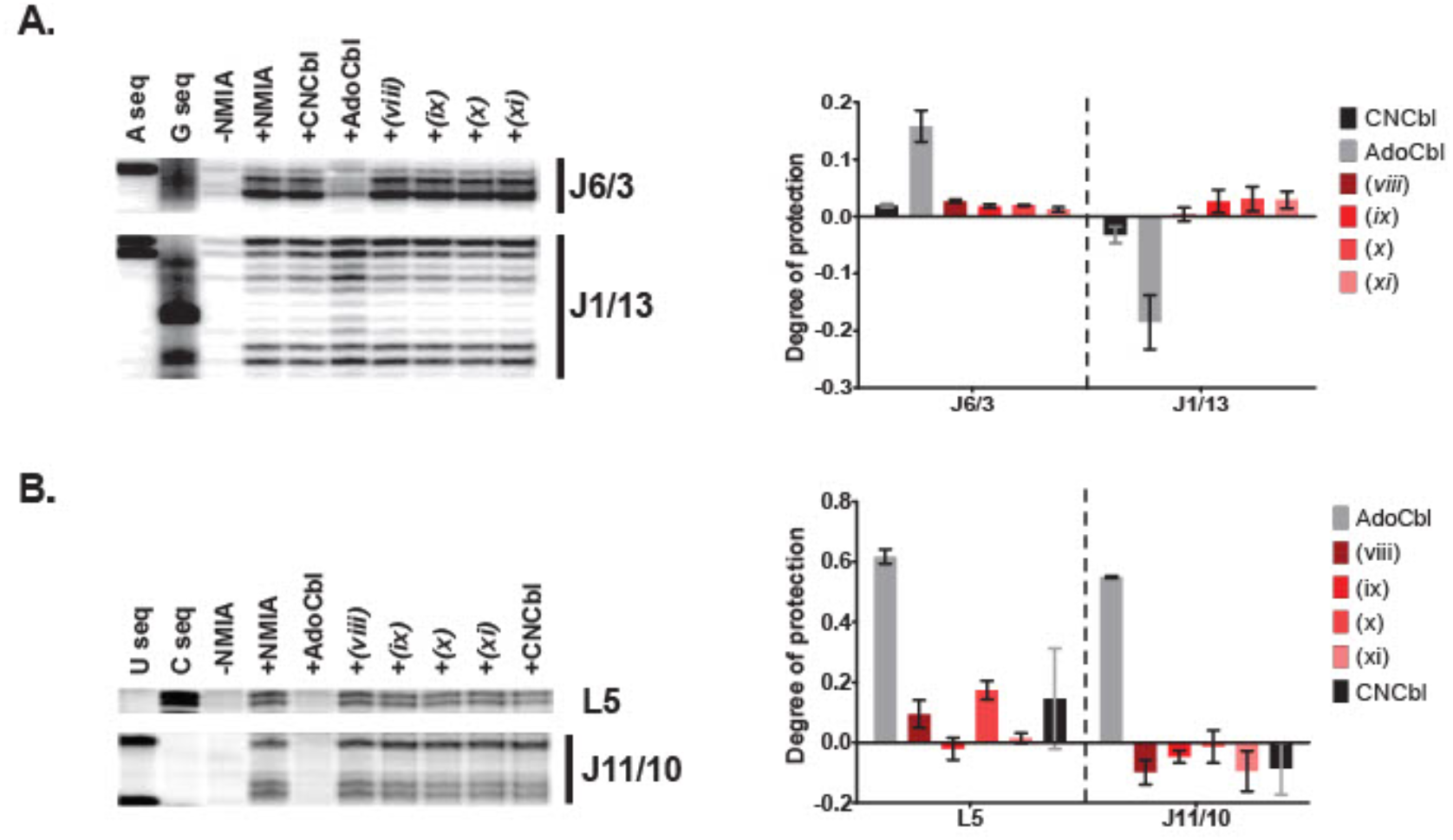
Structural probing by SHAPE of beta-axial Cbl derivatives binding to AdoCbl-selective riboswitches. Representative gels are shown and the results are quantified (n=3, error is represented as SEM). A) *Env50*, J6/3 and J1/13 regions. B) *btuB*, L5 and J11/10 regions.

### Fluorescence displacement to assess binding

As a parallel method to quantify binding of Cbl derivatives to the riboswitches, we used a fluorescent displacement assay. This approach is common for investigating RNA-small molecule interactions and has been used to investigate other riboswitches binding to alternative ligands.^19, 46^ The fluorescent probe used in this assay is a previously characterized fluorophore conjugated form of CNCbl, CNCbl-5xPEG-ATTO590 (**Figure S5A**).^1^ Conjugation of an ATTO 590 fluorophore to the ribose 5’-hydroxyl group of CNCbl via a 5x polyethylene glycol (PEG) linker results in 90% quenching of the fluorophore. However, the conjugated probe binding to *env8* results in 4.9-fold fluorescence induction. Titration of *env8* into CNCbl-5xPEG-ATTO590 gives an apparent equilibrium dissociation constant (K_D_) of 28 ± 7 nM as calculated from the fluorescence read-out (**Figure 4A**), comparable to measurements determined by isothermal titration calorimetry (ITC) in the same conditions (34 ± 9 nM).^1^

**Figure 4.**
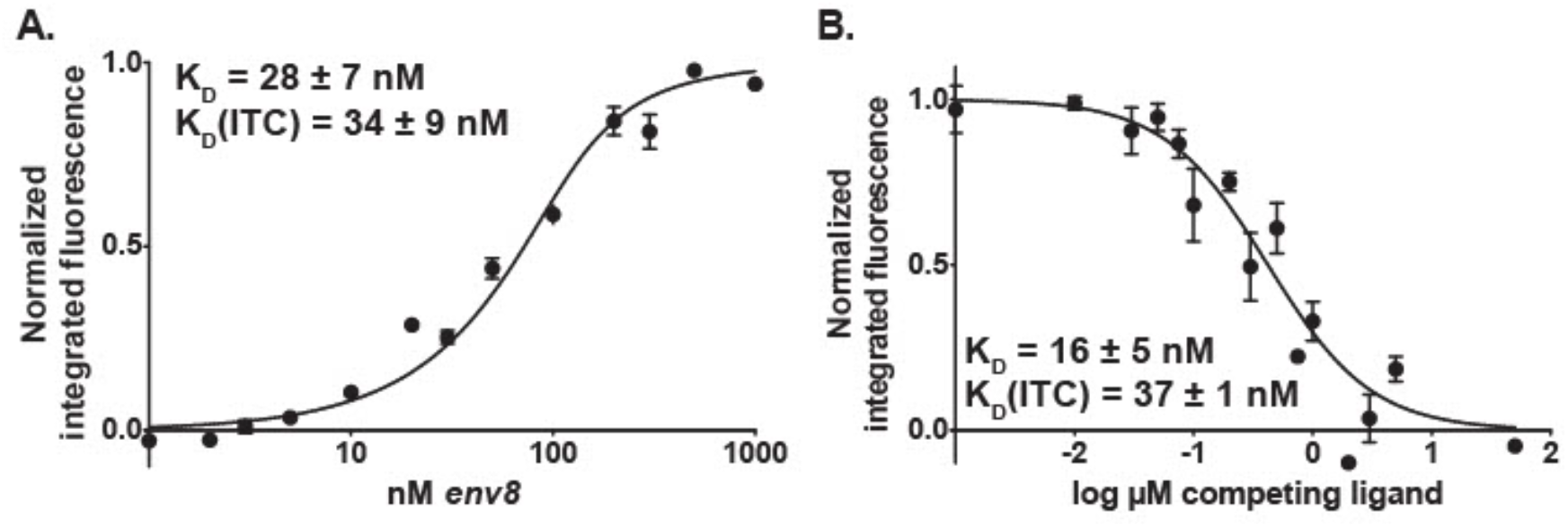
Validation of fluorescence-based affinity measurements. A) Determination of K_D_ by fluorescence induction upon CN-Cbl-5xPEG-ATTO590 probe binding to *env8* (n=9). B) Determination of K_D_ by fluorescence displacement assay, where CNCbl and CN-Cbl-5xPEG-ATTO590 compete for binding to *env8* (n=4). Points represent the average of all replicates and error is represented as SEM. K_D_s determined by ITC are included for comparison.^1^

To quantitatively assess derivative binding, we titrate a Cbl derivative into *env8* saturated with CNCbl-5xPEG-ATTO590, displacing the fluorescent probe. Resultant fluorescence data can be used to determine the concentration of unlabeled competitor that reduces the binding of the probe by half (IC_50_), which can be used to determine K_D_. To validate this method, we titrated a ligand with a known binding affinity for *env8* (CNCbl) into the *env8*-probe complex. The resultant K_D_ of 16 ± 5 nM (**Figure 4B**) is comparable to that determined by ITC (37 ± 1 nM)^1^ and similar to the K_D_ of hydroxocobalamin, another Cbl form with a small beta-axial moiety, as measured by smFRET (5 ± 3 nM).^47^ These data indicate the assay is a reliable method for quantitative assessment of Cbl derivative binding.

### Beta-axial derivatives are comparable to CNCbl in binding *env8*

Using the displacement assay, we surveyed the panel of Cbl derivatives for *env8* binding (**Figure 5, Table 1**). In confirmation of the SHAPE results, every derivative showed some degree of binding and several were highly competitive. There is general agreement that the derivatives that induce a higher degree of SHAPE protection have lower K_D_s. The derivatives that bound most tightly to *env8* relative to CNCbl are beta-axial derivatives *viii* and *ix* which have K_rel_ values of less than 10. Notably, these are the two derivatives that displace A20, as observed by SHAPE. In general, the lactone and *meso* derivatives bound *env8* with lower affinity than most of the beta-axial derivatives with K_rel_ values of 30 and above. Together, these observations indicate that modifications to the beta-axial position of Cbl are tolerated by *env8* better than modifications to the periphery of the corrin ring, although all tested modifications still allow for some level of binding. Further exploration of the chemical space around the beta-axial derivatives may be a productive route to compounds that rival CNCbl in their affinity for class-II Cbl riboswitches.

**Figure 5.**
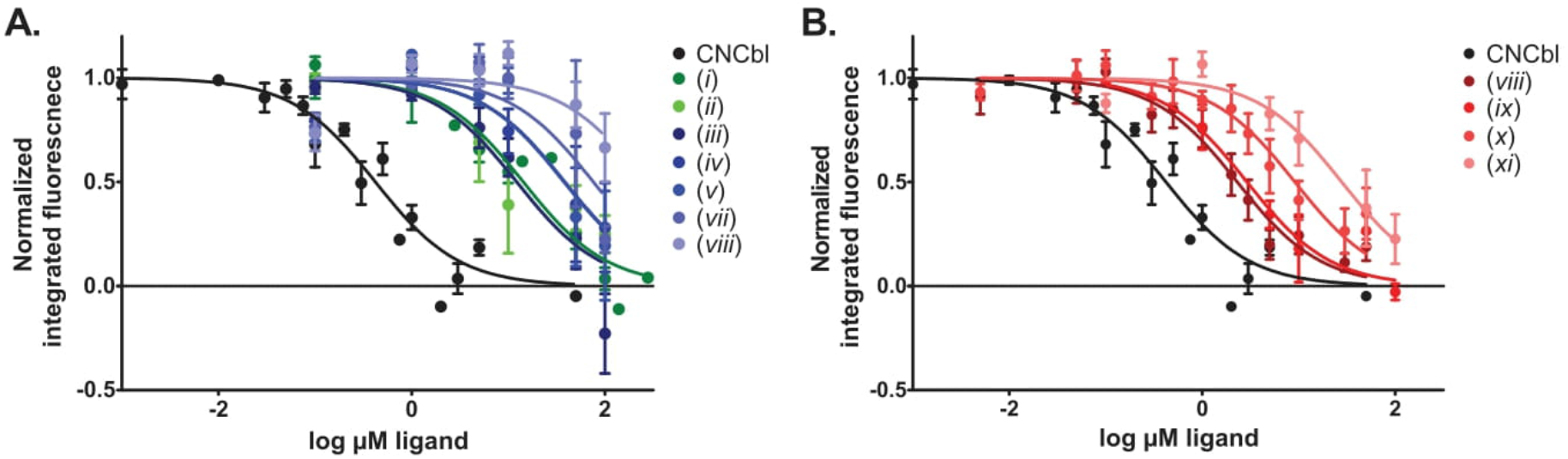
Cbl derivatives binding to *env8*, analyzed by fluorophore displacement assay. A) Lactone and *meso* derivatives. B) Beta-axial derivatives. (n=2-4) Points represent the average of all replicates and error is represented as SEM.

**Table 1.** Binding affinities of Cbl derivatives to *env8* as determined by fluorescence displacement assay.

| Ligand | Derivative Class | $K_D$ (μM)<sup>a</sup> | $K_{rel}$<sup>b</sup> |
| --- | --- | --- | --- |
| CN-Cbl | N/A | $0.016 \pm 0.005$ | 1.0 |
| (i) epi-e-lactone | corrin ring | $0.49 \pm 0.11$ | 30 |
| (ii) c-lactone | corrin ring | $0.72 \pm 0.42$ | 44 |
| (iii) meso-NH<sub>2</sub> | corrin ring | $0.60 \pm 0.35$ | 37 |
| (iv) meso-imine | corrin ring | $0.68 \pm 0.23$ | 42 |
| (v) meso-2°amine | corrin ring | $0.71 \pm 0.04$ | 43 |
| (vi) meso-NO<sub>2</sub> | corrin ring | $2.3 \pm 0.8$ | 142 |
| (vii) meso-amide | corrin ring | $21 \pm 14$ | 1288 |
| (viii) phenyl-NO<sub>2</sub> | beta axial | $0.091 \pm 0.038$ | 5.6 |
| (ix) β-Si | beta axial | $0.12 \pm 0.06$ | 7.2 |
| (x) phenyl-OMe | beta axial | $0.39 \pm 0.15$ | 24 |
| (xi) alkyne-OH | beta axial | $1.5 \pm 1.0$ | 92 |

<sup>a</sup>n=3-4; error reported as SEM
<sup>b</sup> $K_{rel} = K_{D,deriv}/K_{D,CNCbl}$

The results for the beta-axial derivatives can be further interpreted to infer how to successfully target *env8*. The exception to the superior binding of the beta-axial derivatives is *xi* which was the third worst binder with a K_rel_ of 92. This is also the only beta-axial derivative with a methylene bridge proximal to the alkyne in the beta-axial moiety, making it more structurally similar to AdoCbl (**Figure 1A**). This sp^3^-hybridization likely projects the aromatic ring towards J6/3, consistent with the lack of increased SHAPE reactivity of A20 upon *xi* binding. If *xi* projects into the same segment of the purine stack as AdoCbl, this also likely causes its reduced binding affinity. The other beta-axial derivative that does not displace A20 is derivative *x*, although it is most structurally similar to *viii*. We hypothesize that this is due to the difference between the OMe versus NO_2_ terminal groups. The nitro group in derivative *viii* withdraws electrons from the phenyl ring, which enhances π-stacking interactions.^48^ Thus, *viii* may help stabilize the purine stack after A20 displacement. In contrast, the methoxy group in *x* is an electron donating group which increases electronic repulsion during base stacking,^48^ disfavoring the replacement of A20 with *x* in the purine stack. This hypothesis is supported by the reduced affinity of *x*, which is about four times worse than *viii*. The most surprising result of the beta-axial derivatives is *ix*, which has productive binding through displacement of A20 despite not possessing an aromatic ring to intercalate into the purine stack, indicating that π-stacking is not essential for facilitating displacement of A20.

### Fluorescence displacement reveals binding undetected by SHAPE

For fluorescence displacement assays with *env50* and *btuB*, we synthesized an additional Cbl-fluorophore conjugate. Derivative *viii* was modified in the same manner as CNCbl-5xPEG-ATTO590, resulting in a PhNO_2_-Cbl-5xPEG-ATTO590 probe (**Figure S5B**). This allowed for direct analysis of binding to this Cbl derivative. Measurement of fluorescence induction upon binding showed *env50* has slightly higher affinity for the PhNO_2_ probe than the CN probe (**Figure 6A**). The observed K_D_ for the CN probe is consistent with a previously reported K_D_ for MeCbl as measured by ITC.^24^ However, using either probe with *env50* to quantitatively determine derivative K_D_ values requires saturation of the RNA. Given the high nM affinity of these probes for *env50*, large amounts of each derivative would be necessary to fully define a binding curve. Additionally, we observed that high concentrations of derivative decreased the background fluorescence from either quenching of the probe or the inner filter effect, complicating quantification of the data.

**Figure 6.**
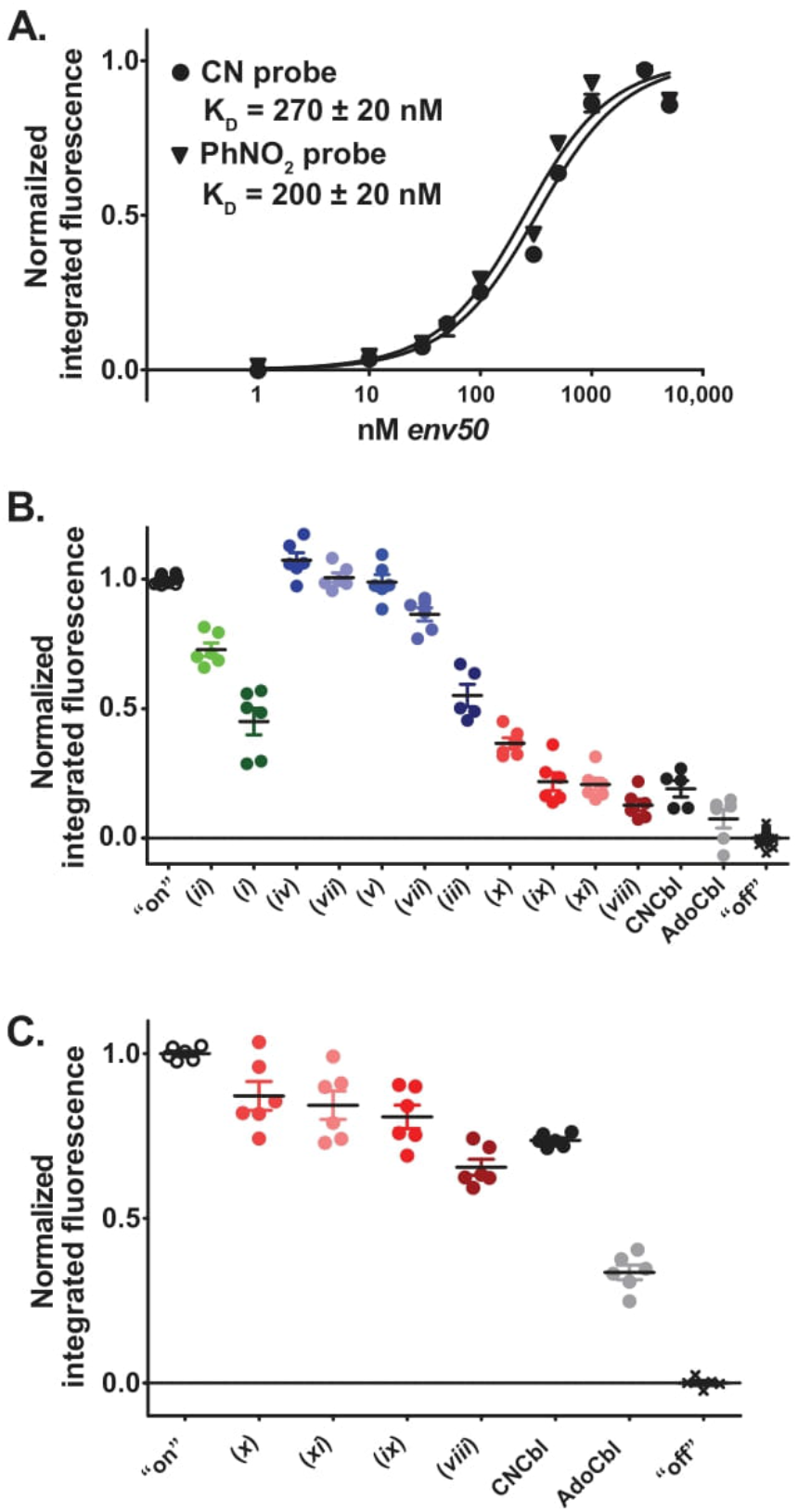
Cbl derivatives binding to *env50*, analyzed by fluorescence assays. A) Determination of K_D_ by fluorescence induction upon CN-Cbl-5xPEG-ATTO590 and PhNO_2_-Cbl-5xPEG-ATTO590 probes binding to *env50* riboswitch (n=3). Points represent the average of all replicates and error is represented as SEM. B) Binding of Cbl derivatives (1 µM) to *env50* by fluorescence displacement assay (n=3). Values are normalized so “on” = probe fluorescence bound to *env50* without competing ligand and “off” = probe fluorescence unbound to RNA. C) The same as B but competing ligands are at 100 nM. Error is represented as SEM.

Given the above complication, we changed the assay conditions to qualitatively assess *env50* binding to the Cbl derivatives. Binding was evaluated at a 1:1 probe:derivative ratio for weakly binding compounds and a 10:1 probe:derivative ratio for high affinity derivatives (**Figure 6B**). The Cbl derivatives displayed a range of probe displacement. The observed trend in probe displacement is similar to that for *env8* with three exceptions discussed below. As expected, the beta-axial derivatives were the best competitors while the *meso* and lactone derivatives displayed poor to moderate competition. CNCbl was the second most effective competitor, although derivatives *ix* and *xi* performed nearly as well. Derivative *viii* outcompeted CNCbl, consistent with the above direct binding data. However, derivative *viii* was not as effective a competitor as AdoCbl, reinforcing *env50*’s preference for this Cbl form. Using the beta-axial derivatives to compete at lower concentrations than the probe confirmed the superior binding capability of AdoCbl (**Figure 6C**).

Three derivatives do not follow the same trend in affinities between *env8* and *env50*. Two lactone derivatives, *i* and *ii*, which were similar to the *meso* derivatives in affinity for *env8*, appear to have improved affinity for *env50*. Additionally, derivative *xi* shows marked improvement in binding *env50* compared to *env8*; *xi* ranked with the worst of the binders for *env8*, while it ranks among the best for *env50*. We hypothesize that *xi*’s structural similarity to AdoCbl may allow for better binding to AdoCbl-selective riboswitches like *env50*.

The fluorescence displacement results contrast the *env50* SHAPE data, which did not indicate binding for either CNCbl or any of the Cbl derivatives. One potential explanation for these results is a reaction between the SHAPE chemical probing agent and Cbl that inactivates binding. However, incubation of CNCbl with the probing agent did not alter its binding to *env50* (**Figure S6**). The negative SHAPE results may also be due to reaction conditions, indicative of a high k_off_ rate of the derivatives, or the higher K_D_s may present a limitation for the methodology. Evidence suggests that SHAPE-based screening may be biased towards high affinity ligands, although K_D_s up to hundreds of micromolar have been detected.^49^ Disagreement between the results of SHAPE and another method has been seen previously; the *Tte*-PreQ_1_ riboswitch did not show an altered SHAPE signature for synthetic ligand binding despite a change in RNA flexibility suggested by molecular dynamics simulations.^50^ The inconsistency between our results suggests that SHAPE may not be ideal for screening weakly binding compounds during initial phases of discovery of an RNA targeting compound. These observations highlight the utility of parallel approaches to assess binding in novel systems.

To examine binding of our panel of Cbl derivatives to *btuB*, we first assessed direct binding of the CN and PhNO_2_ probes. The highest tested concentration of RNA (10 µM) did not induce fluorescence in the probes, consistent with the SHAPE results (**Figure S7**). Together the *btuB* fluorescence and SHAPE data suggest that *btuB* binding is highly specific for AdoCbl. Due to the lack of binding detected with either probe, we could not use the displacement assay to determine binding affinities for the Cbl derivatives. In light of these and the SHAPE results, we do not expect the derivatives to productively bind the class-I Cbl riboswitches. Targeting AdoCbl-specific riboswitches likely requires development of stable beta axial moieties that better mimic the Ado moiety.

### *Env8*’s binding pocket is flexible but largely unalterable for beta-axial derivative binding

To determine whether observed plasticity of the J3/4-J6/3 purine spine is specific to adenosine, we examined the role of A20 in Cbl binding. This position was mutated to each of the other nucleotides, and the mutant sequences’ ability to bind CNCbl, *viii*, and *ix* was examined using SHAPE (**Figure 7A**). Despite this position being a purine across the Cbl riboswitch family, all mutants exhibit the same patterns of SHAPE reactivity in the presence of the tested Cbls, including the signature for displacement of nucleotide 20. Thus, the conformational flexibility of this nucleotide is independent of its identity.

**Figure 7.**
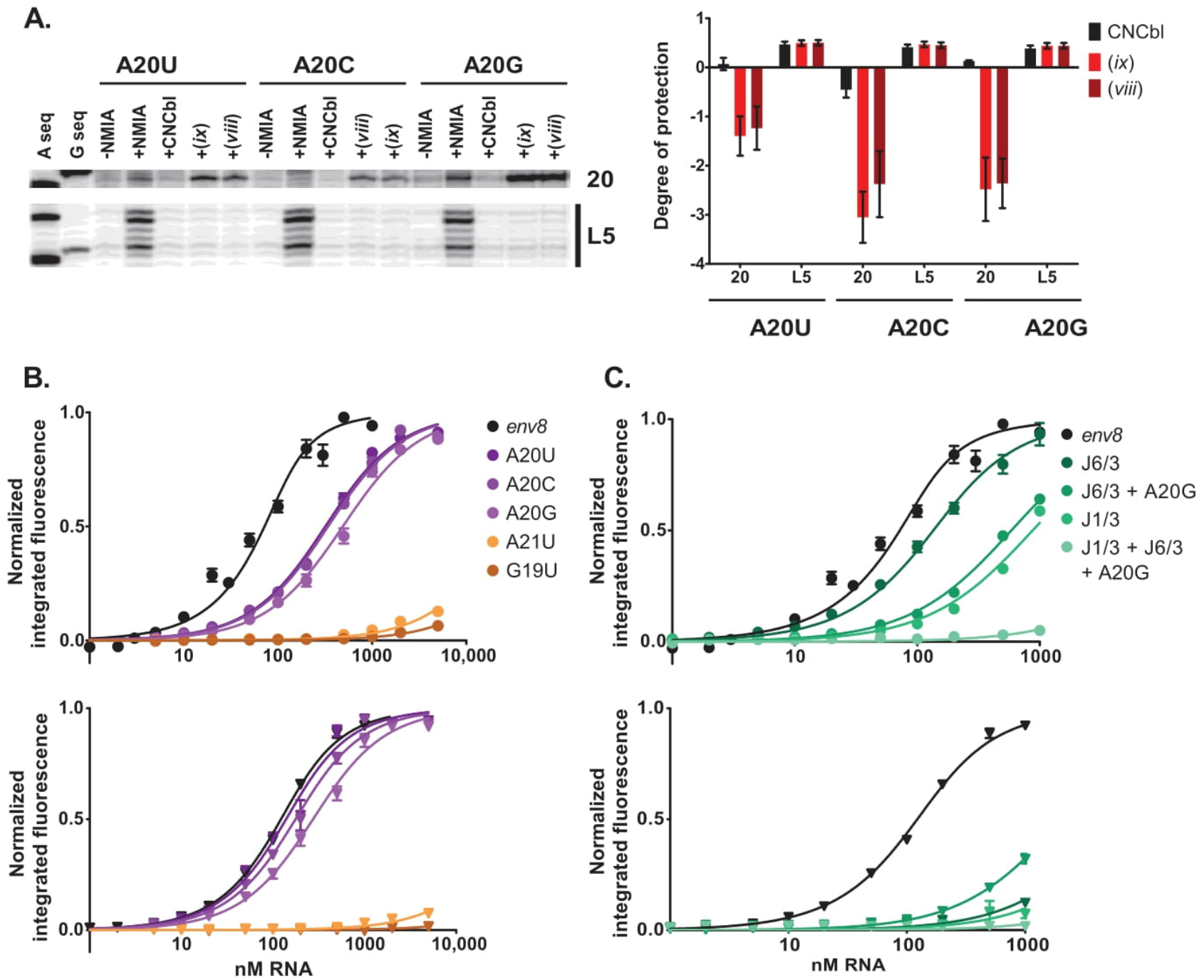
*Env8* binding pocket mutants binding to Cbl derivatives. A) A20 mutants were analyzed for binding to CNCbl, *viii*, and *ix* by SHAPE. Nucleotide A20 and region L5 are shown in a representative gel and are quantified (n=2). B) Determination of K_D_ by fluorescence induction upon CN-Cbl-5xPEG-ATTO590 (top) and PhNO_2_-Cbl-5xPEG-ATTO590 (bottom) binding to *env8* A20 mutants. C) Same as B, but mutations to regions J6/3 (A68G) and J1/3 (ΔA9, G10U, G12A), alone and in combination with A20G. (n=3-4) Error is represented as SEM.

Since pyrimidines also support Cbl binding, we hypothesized that they may be easier to displace and thus promote PhNO_2_ binding. We quantified the binding of the CN and PhNO_2_ probes to the above mutants along with mutations in the flanking positions (G19U and A21U). Mutation of positions flanking nucleotide 20 do not support binding, indicating the importance of these residues (**Figure 7B, Table 2**). The position 20 mutants are less detrimental to the PhNO_2_ probe binding than the CN probe. Notably, each of the A20 mutants prefers binding the PhNO_2_ probe over the CN probe.

**Table 2.** Binding affinities of CN-Cbl-5xPEG-ATTO590 and PhNO_2_-Cbl-5xPEG-ATTO590 probes to *env8* variants.

| ***env8* variant** | **CN probe K<sub>D</sub>**<br>**(nM)<sup>a</sup>** | **PhNO<sub>2</sub> probe K<sub>D</sub>**<br>**(nM)<sup>a</sup>** | **K<sub>rel</sub><sup>b</sup>** |
| --- | --- | --- | --- |
| Wild type | 28 ± 7<sup>b</sup> | 73 ± 8 | 0.4 |
| A20U | 270 ± 20 | 90 ± 2 | 3.0 |
| A20C | 290 ± 20 | 130 ± 10 | 2.2 |
| A20G | 450 ± 70 | 230 ± 50 | 2.0 |
| G19U | ND<sup>c</sup> | ND |  |
| A21U | ND | ND |  |
| J6/3 | 90 ± 6 | ND |  |
| J6/3 + A20G | 540 ± 10 | ND |  |
| J1/3 | 820 ± 30 | ND |  |
| J1/3 + J6/3 + A20G | ND | ND |  |

<sup>a</sup> n=3-4; error reported as SEM
<sup>b</sup> K<sub>rel</sub> = K<sub>D,CN</sub>/K<sub>D,PhNO2</sub>
<sup>c</sup> ND: binding not detectable.

We next sought to assess whether nucleotides supporting the binding pocket influence the plasticity of nucleotide 20.^24^ A series of mutations were made to nucleotides in regions J6/3 and J1/3, alone and in combination with the A20G mutation. Given that *env50* shows reduced affinity for the beta-axial derivatives compared to *env8*, we mutated the *env8* regions to match *env50* to test our hypothesis. Within this set of mutations, binding to the CN probe is progressively diminished, recapitulating prior observations (**Figure 7C, Table 2**).^24^ In stark contrast, mutations beyond A20G that support CNCbl binding are severely deleterious to PhNO_2_ binding. This reveals the flexibility of nucleotide 20 is dependent upon the surrounding RNA architecture which suggests that accommodation of beta-axial derivatives may be restricted to a subset of class-II riboswitches. In the development of an FMN riboswitch targeting compound, a similar phenomenon was observed in which the lead compound was highly specific for the *Clostridium difficile* FMN riboswitch.^17^ This property enabled the lead compound to target *C. difficile* in an animal model while leaving the remainder of the microbiome largely unaffected—a potentially important feature of a desired antimicrobial agent.^17, 51^

### The beta-axial Cbl derivatives can drive biological function

The above results demonstrate that a set of beta-axial Cbl derivatives productively bind the *env8* riboswitch, but binding *in vitro* may not drive regulatory function in cells. Regulatory function may not occur for several reasons, including an inability to import, an inability to bind the riboswitch within a relevant timescale, or an inability to occlude the ribosome binding site to repress translation.^40^ To determine whether the derivatives repress gene expression, we tested the beta-axial Cbl derivatives in a previously established cell-based reporter assay in *E. coli*.^40^ In this assay, the *env8* riboswitch is upstream of a fluorescent protein (FP) reporter whose expression is repressed by Cbl-dependent occlusion of the ribosome binding site. The beta-axial derivatives that productively bind *env8 in vitro* also repress FP expression (**Figure 8**). While the repressive activity of these ligands was less than that of CNCbl, they generally agree with their binding affinities. Derivative *vii*, the worst binder *in vitro*, does not show this repressive effect, indicating that biological activity of the other derivatives is due to riboswitch binding. Thus, these derivatives serve as outstanding lead candidates for further development as inhibitors of genes involved in Cbl metabolism that are regulated by class-II Cbl riboswitches.

**Figure 8.**
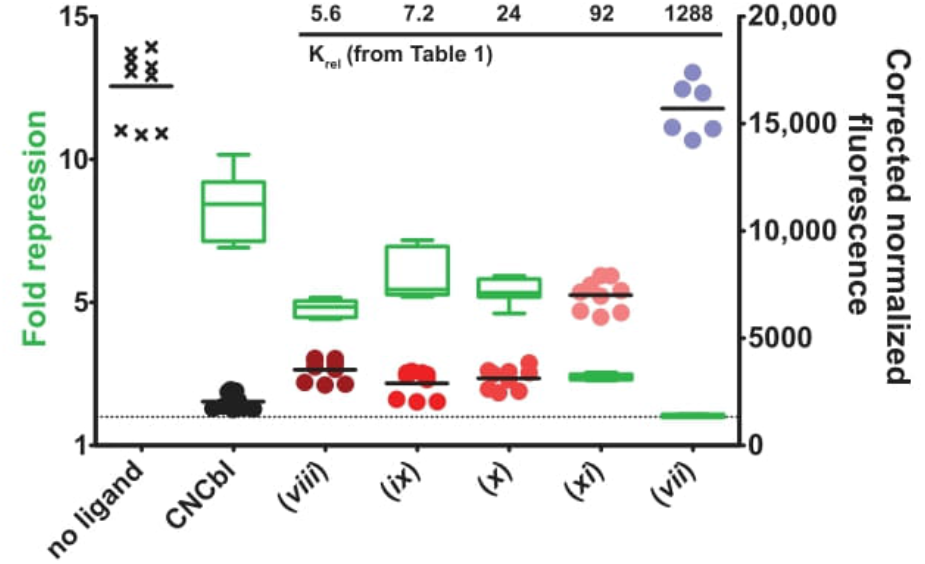
*In vivo* fluorescence reporter assay shows Cbl derivatives are capable of regulating translation of a protein (mNeonGreen) under control of the Cbl riboswitch (n=3). Protein fluorescence is measured on the right y-axis and shown with symbols. Fluorescence fold repression compared to the no ligand condition is measured on the left y-axis and shown with green box-and-whisker plots. *vii*, shown to have low affinity for *env8* by displacement assay (Figure 3), was used as a negative control. K_rel_ values of the Cbl derivatives from Table 1 are included for reference.

### Summary and Outlook

In this work, we have demonstrated that class-II Cbl riboswitches are able to bind photostable beta-axial cobalamin derivatives and regulate gene expression in *E. coli*. Chemical probing revealed that a critical aspect of recognition of the beta-axial derivatives is displacement of a nucleotide from a purine stack central to the binding pocket. This displacement mechanism is likely supported through replacement of the displaced nucleotide by intercalation of the derivative functional group as has been seen in other RNAs.^36^ Thus, this study adds to a growing body of evidence that exploiting RNA’s structural plasticity is a productive path in drug discovery efforts.

Complementary to the class-II Cbl riboswitch findings is the observation that an increase in AdoCbl selectivity, integral to class-I Cbl riboswitches, appears to be deleterious to plasticity. It has been hypothesized that class-I Cbl riboswitches are incapable of discriminating between AdoCbl and similar antivitamin B_12_ forms and some evidence supports this.^25, 27, 52^ However, we show that the *E. coli btuB* Cbl riboswitch did not bind any of the Cbl derivatives tested through either SHAPE or fluorescent methods. This suggests that displacement of the equivalent A20 position in this riboswitch is disallowed, which may be a result of the plasticity of the nucleotide on the opposite strand in J6/3 that is required to accommodate the Ado. Structural analysis of the novel binding modes discovered here will reveal how the beta-axial derivatives specifically interact with the Cbl riboswitch binding core and yield potential insights into how differences between class-I and class-II result in binding specificities. Thus, this work provides new insights to target a broadly distributed RNA in bacteria that regulates metabolism of an essential cofactor for survival and virulence in a number of medically important pathogens.

## Materials and Methods

### RNA synthesis and preparation

All RNAs used in this study (sequences in **Table S1**) were made using dsDNA templates amplified by PCR,^53^ transcribed with T7 RNA polymerase, and purified using denaturing PAGE.^53^ Purified RNA was buffer exchanged and concentrated into milli-Q H_2_O using centrifugal concentrators (Amicon). Final RNA concentrations were calculated using A_260_ and molar extinction coefficients determined from the summation of the individual bases.

### Cobalamin derivative and probe synthesis

Derivatives *i-xi* and the CN probe were synthesized according to previously described procedures.^1, 29, 32, 54-55^ See Supporting Information for further details and synthesis of the PhNO_2_ probe.

### Fluorescence induction direct binding assay

RNAs were titrated into 30 µL reactions in a Corning 384-well plate such that the final concentrations were 100 nM CNCbl-5xPEG-ATTO590 and 1x of RNA buffer (100 mM KCl, 10 mM NaCl, 1 mM MgCl_2_, 50 mM HEPES, pH 8.0).^1^ The concentration of probe could not be substantially lowered due to the sensitivity of the plate reader. Reactions were done in at least triplicate. Reactions were incubated for at least 30 minutes at room temperature in the dark before reading. Time to equilibration was experimentally validated (**Figure S8**). ATTO 590 fluorescence was monitored (594 nm excitation, 620-670 nm emission) using a BMG Labtech CLARIOstarPLUS microplate reader. Fluorescence values were background corrected by subtracting buffer fluorescence values, integrated over all wavelengths, and normalized to the average integrated fluorescence of the no RNA control reactions. The corrected normalized integrated fluorescence values were plotted versus log(nM RNA) in GraphPad Prism. Technical replicates were fit to the quadratic binding equation with one transition,

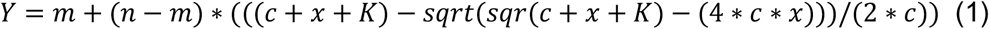

where *Y* is the corrected normalized fluorescence value, *m* is the lower baseline, *n* is the upper baseline, *c* is the probe concentration, *x* is RNA concentration, and *K* the K_D_. The quadratic equation was used due to the K_D_ being lower than the concentration of the probe.^56^ Calculated K_D_s are the average of multiple biological replicates, and SEM was calculated from those average K_D_s. Biological replicates were combined for the final graphs shown in the figures.

### Fluorescence displacement assays

In experiments using *env8*, competing ligands were titrated into 30 µL reactions in a Corning 384-well plate such that the final concentrations were 1 µM CNCbl-5xPEG-ATTO590, 100 nM RNA, and 1x of RNA buffer. Reactions were done in technical duplicates or triplicates and in 2-6 biological replicates. Equilibration, plate reading, background subtraction, and fluorescence value integration were done as above. Values were plotted as µM competitor ligand versus integrated fluorescence in GraphPad Prism. Technical replicates were fit to a nonlinear regression curve with the HillSlope parameter set equal to 1.0 and normalized. Fitting gave an IC_50_ which was used to determine K_D_ using the equation

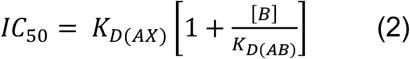

where *A* is the RNA, *B* is CNCbl-5xPEG-ATTO590, and *X* is the competing ligand.^57^ A K_D_ of 34 nM between *env8* and CNCbl-5xPEG-ATTO590 was used.^1^ Biological replicates consisting of normalized technical replicates were combined for the final graphs shown in the figures. They were refit to a nonlinear regression curve with the HillSlope set equal to 1.0, the bottom set equal to 1.0, and the top set equal to 0. Calculated K_D_s are the average of multiple biological replicates, and SEM was calculated from those average K_D_s.

For qualitative analysis, 30 µL reactions had final concentrations of 1 µM CNCbl-5xPEG-ATTO590, 100 nM RNA, 1x of RNA buffer, and either 1 µM or 100 nM of competing ligand. Corrected and integrated fluorescence values were normalized to those of wells with no competing ligand (“on” = 1) and with no RNA (“off” = 0) and were plotted in GraphPad Prism.

### Selective 2’-Hydroxyl Acylation Analyzed by Primer Extension (SHAPE)

Chemical structure probing was performed using *N*-methylisatoic anhydride (NMIA) as previously described with modifications.^39^ See Supporting Information for details.

### Cell-based reporter assays

Conducted as previously reported^24^ with the following changes: 0.5 μL of the overnight culture was added to 500 μL of medium and 150 μL were added to wells in a Costar 96-well plate.

## Supporting information

Document with all Supplemental Information

## Supporting Information

Figures S1-8: *env8* sequence secondary structure, full SHAPE gels and additional quantification for *env8, env50*, and *btuB*, chemical structures of probes, chemical probing agent control binding assay, fluorescence turn-on upon probes binding to *btuB*, and binding equilibration assays; Tables S1-2: RNA and DNA primer sequences and degree of protection values from SHAPE gels; additional methods including characterization of synthesized cobalamin compounds; supplemental references (DOC). Compound characterization checklist (XLS).

## Acknowledgement

The authors wish to acknowledge and thank Esther Braselmann for her contribution to the initial development of this work in choosing cobalamin derivatives to include on the panel and conducting preliminary binding assays.

## Funding Sources

NIH R01 GM133184

NIH R01 GM073850

NIH T32 GM065103

Polish National Agency for Academic Exchange (no. PPN/BEK/2020/1/00219/U/00001)

Foundation for Polish Science (START scholarship no. 092.2021)

National Science Centre, Poland (MAESTRO 2020/38/A/ST4/00185)

## Author Contributions

SRL, AEP, and RTB conceptualized and designed the study. AJW synthesized and characterized the cobalamin compounds. SHS contributed to the data in Figure 5. SRL conducted all experiments and analyzed all data, with input from all authors. SRL and RTB wrote the manuscript with edits from all authors.

