## Supplementary material for "Targeting riboswitches with beta-axial substituted cobalamins": Document with all Supplemental Information

**Supporting Information**

Shelby R. Lennon[1], Aleksandra J. Wierzba[1,2], Shea H. Siwik[1], Dorota Gryko[3], Amy E. Palmer[1,2] and Robert T. Batey[1,*]

[1]Department of Biochemistry, University of Colorado, Boulder, CO 80309-0596, USA

[2]BioFrontiers Institute, University of Colorado, Boulder, CO 80303 – 0596, USA

[3]Institute of Organic Chemistry, Polish Academy of Sciences, Kasprzaka 44/52, 01-224 Warsaw, Poland

**Table of Contents:**

**Figure S1:** env8 secondary structure P. 3

**Figure S2:** SHAPE gels of *env*8 bound to lactone/*meso* derivatives P. 4

**Figure S3:** SHAPE gels of env50 bound to beta axial derivatives P. 5

**Figure S4:** SHAPE gels of *btuB* bound to beta axial derivatives P. 6

**Figure S5:** Chemical structures of cobalamin probes P. 7

**Figure S6:** Control of NMIA reactivity with cobalamin P. 8

**Figure S7:** Analysis of cobalamin probes binding to *btuB* P. 9

**Figure S8:** Empirical determination of time to equilibration P. 10

**Table S1:** Sequences used in this study P. 11

**Table S2:** Raw quantification of SHAPE gels P. 13

**Supplemental Methods** P. 16

**Supplemental References** P. 25

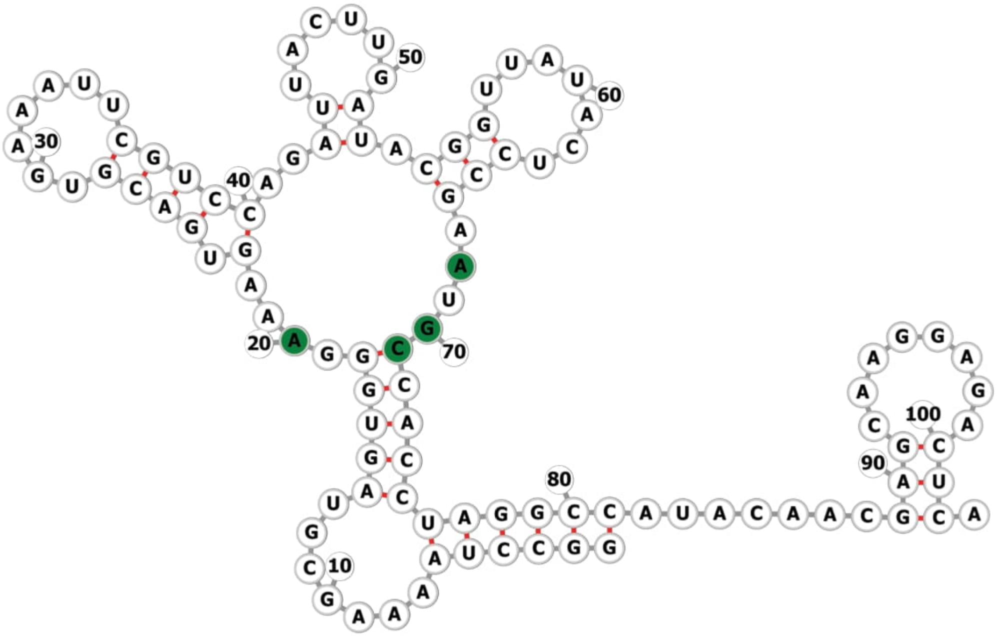

**Figure S1.** *env8* secondary structure. Nucleotides that clash with Cbl derivatives as seen by molecular docking are colored in green. Image made with forna.[1]

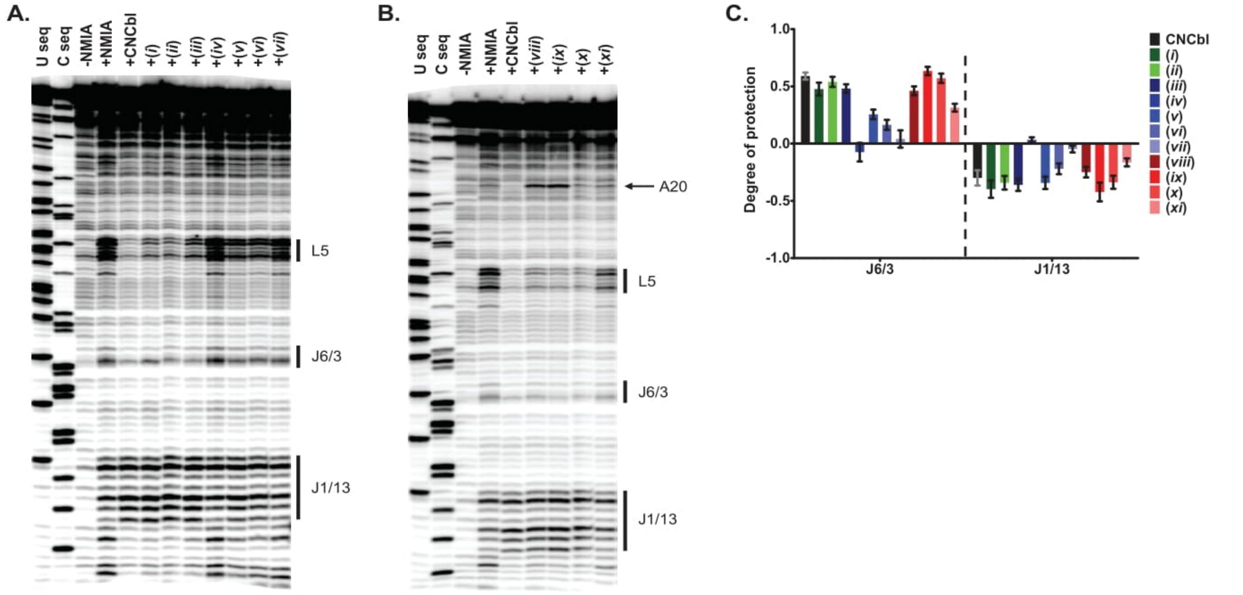

**Figure S2.** Full representative SHAPE gels of *env8* and lactone/*meso* (A) and beta-axial (B) Cbl derivatives, from Figure 2, with key RNA regions marked on the right-hand side. Gels have been straightened and had contrast adjusted by semi-automated footprinting analysis (SAFA) software. C) Quantification of J6/3 and J1/13 regions. n = 6-10, error is represented as SEM.

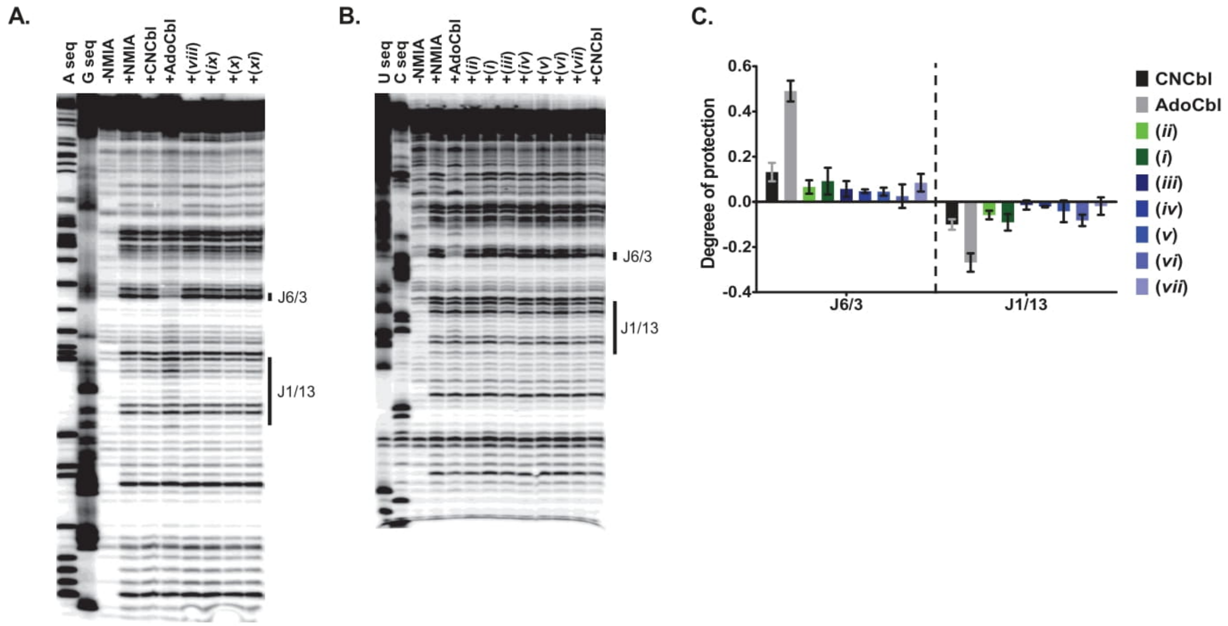

**Figure S3.** Full representative SHAPE gels of *env50* and beta-axial Cbl derivatives (A, from Figure 3) and lactone/*meso* Cbl derivatives (B), with key RNA regions marked on the right-hand side. Gels have been straightened and had contrast adjusted by SAFA. C) Quantification of J6/3 and J1/13 regions for lactone and *meso* derivatives, n = 3, error represents SEM.

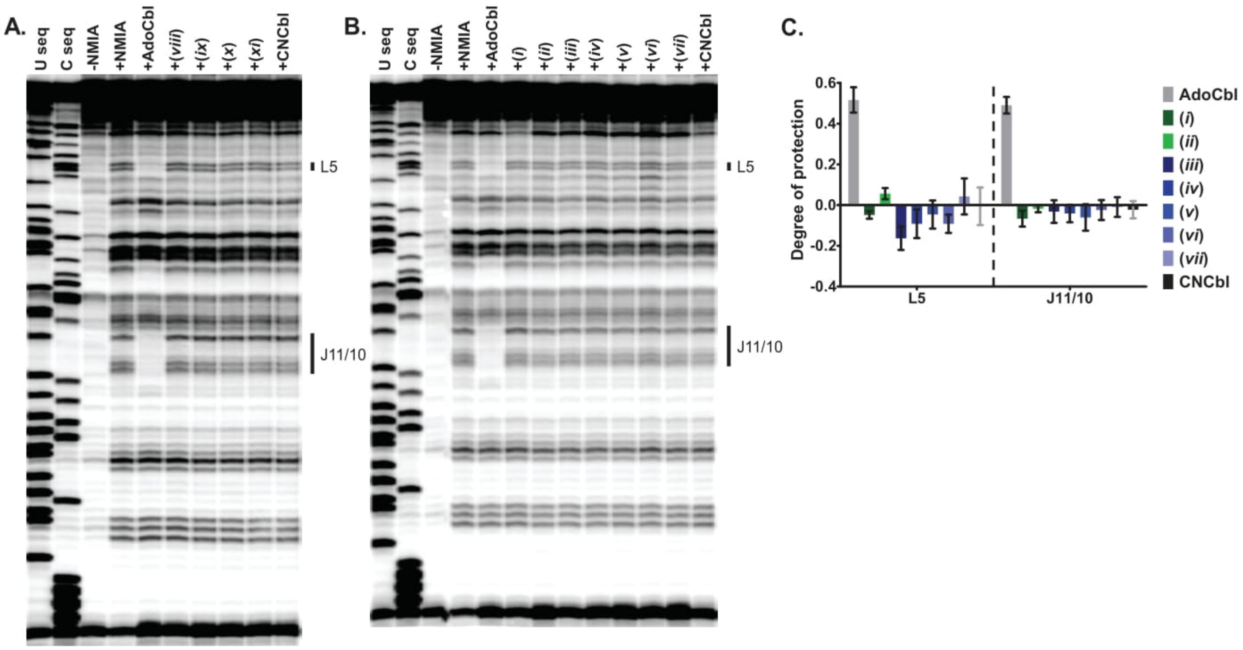

**Figure S4.** Full SHAPE gels of *btuB* and beta-axial Cbl derivatives (A, from Figure 3) and lactone/*meso* Cbl derivatives (B), with key RNA regions marked on the right-hand side. A primer internal to the riboswitch sequence was used for reverse transcription to improve resolution of the 5'-end. Gels have been straightened and had contrast adjusted by SAFA. C) Quantification of L5 and J11/10 regions for lactone and *meso* derivatives, n = 3, error represents SEM.

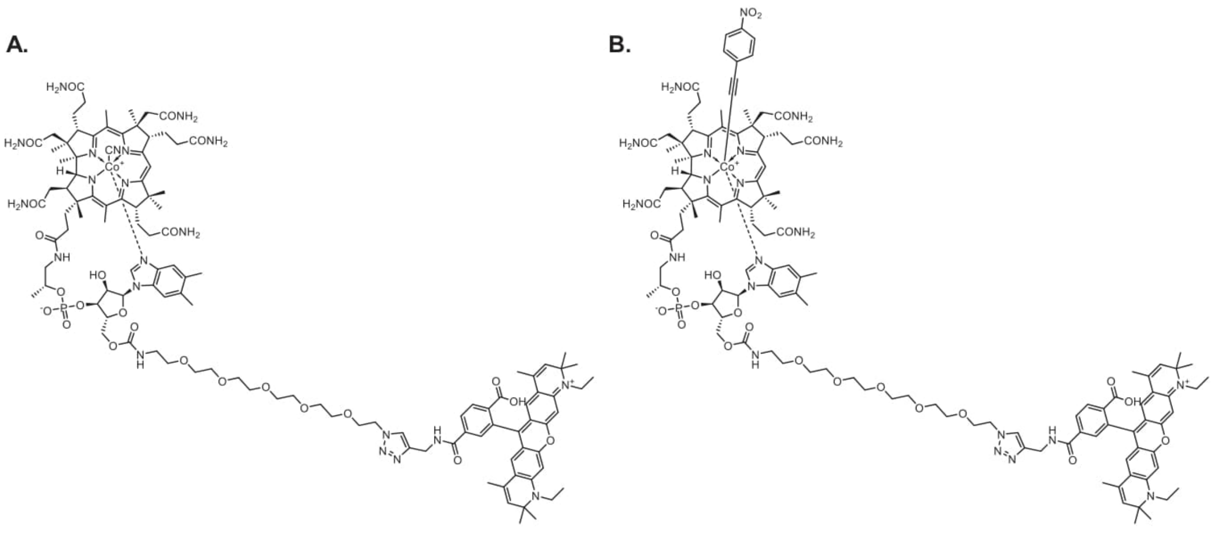

**Figure S5.** Chemical structures of the (A) CNCbl-5xPEG-ATTO590 and (B) $PhNO_2$-Cbl-5xPEG-ATTO590 probes.

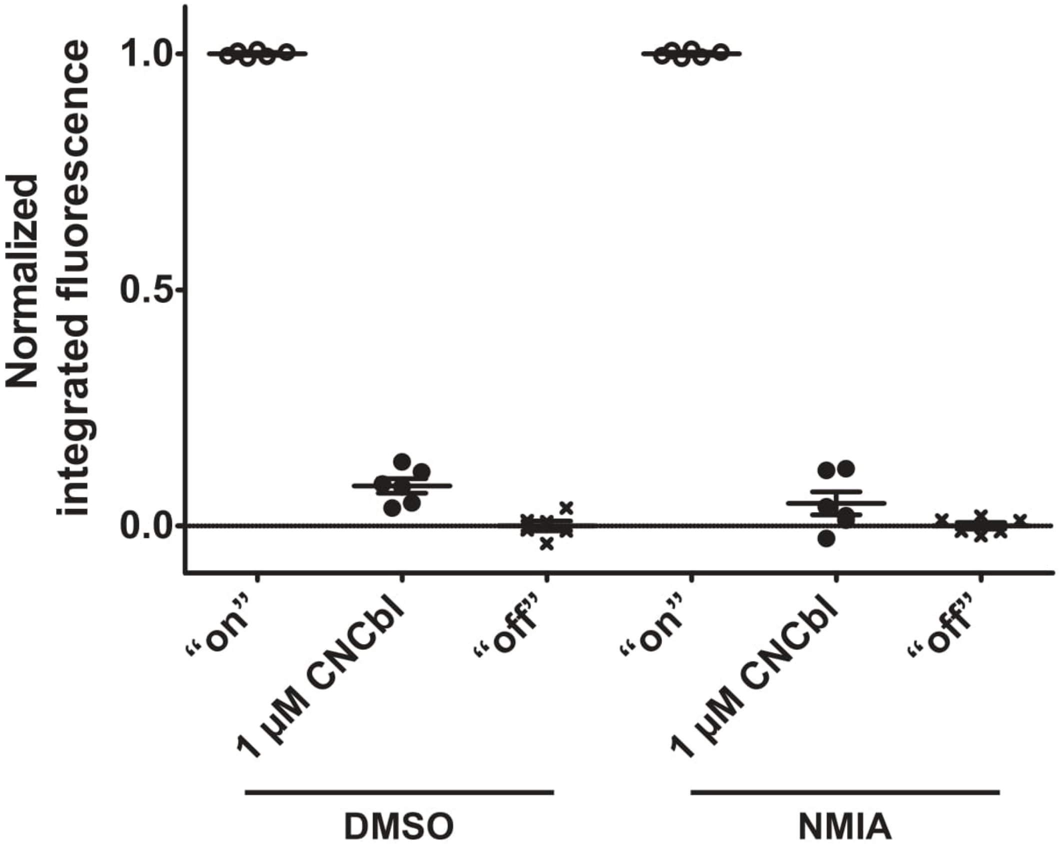

**Figure S6.** CNCbl does not react with SHAPE chemical probe NMIA in a manner that prevents binding to *env50*. CNCbl, incubated in either DMSO (control) or NMIA in conditions used for SHAPE reactions, was used in the fluorescence displacement assay to compete with the CN probe for binding to *env50*. Values are normalized as in Figure 6. n = 3, error represents SEM.

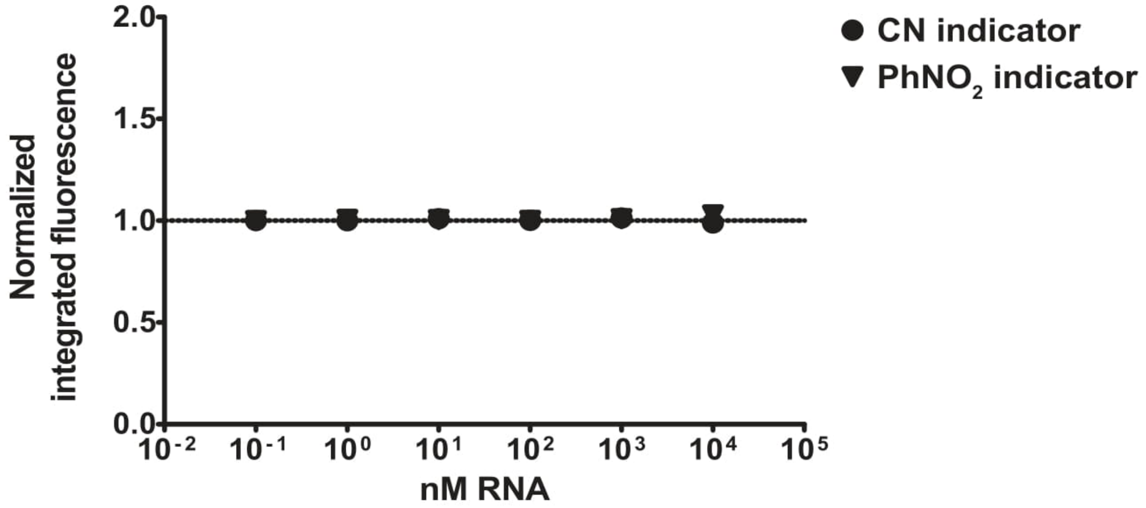

**Figure S7.** Analysis of CN-Cbl-5xPEG-ATTO590 and $PhNO_2$-Cbl-5xPEG-ATTO590 probes binding to *btuB* riboswitch by fluorescence induction (n=3). Error bars, representing SEM, are smaller than symbols.

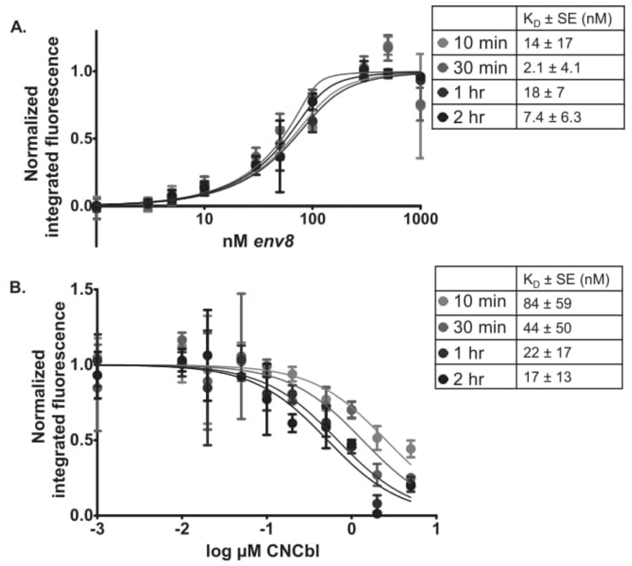

**Figure S8.** Empirical determination of time to equilibration for fluorescence assays. A) Fluorescence induction assay completed as in Figure 4A with equilibration times listed. B) Fluorescence displacement assay completed as in Figure 4B with equilibration times listed. Errors represent SEM of two technical replicates.

**Table S1.** Sequences used in this study.

| Name | Sequence | Genbank accession |
|---|---|---|
| *env8* | GGC CUA AAA GCG UAG UGG GAA AGU GAC GUG AAA UUC GUC CAG AUU ACU UGA UAC GGU UAU ACU CCG AAU GCC ACC UAG GCC AUA CAA CGA GCA AGG AGA CUC | AACY021350931.1 |
| *env50* | GGU ACU GAA UCA UGG UGG GGA ACA AUG UGA AAU UCA UUG ACU GUU CCU GCA ACG GUA AAA GUA AAA UUG AGU CCG AGU GCC ACC CAG UAA AGU CCG CUG UGA GUG AAG GCC AGG AAA AGU CUA ACU C | AACY023404808.1 |
| *E. coli btuB* Cbl riboswitch | GGC CGG UCC UGU GAG GUU AAU AGG GAA UCC AGU GCG AAU CUG GAG CUG ACG CGC AGC UGU UAG GAA AGG UGC CAU GAU GUC GUU AUG CGG ACA CUC GCC AUU CGG UGG GAA GUC AUC AUG UCU UAG UAU CUU AGA UAC CCC UCC AAG CCA GGA GAC CUG CCG GCC AAC GUC GCA UCU GGU UCU CAU CAU CGC GUA AUA UUG AUG AAA CCU GCG GCA | M10112.1 |
| *env8*_SHAPE | GGC CUU CGG GCC AAG GCC UAA AAG CGU AGU GGG AAA GUG ACG UGA AAU UCG UCC AGA UUA CUU GAU ACG GUU AUA CUC CGA AUG CCA CCU AGG CCA UAC AAC GAG CAA GGA GAC UCU CGA UCC GGU UCG CCG GAU CCA AAU CGG GCU UCG GUC CGG UUC | |
| *env50*_SHAPE | GGU ACU GAA UCA UGG UGG GGA ACA AUG UGA AAU UCA UUG ACU GUU CCU GCA ACG GUA AAA GUA AAA UUG AGU CCG AGU GCC ACC CAG UAA AGU CCG CUG UGA GUG AAG GCC AGG AAA AGU CUA ACU CAA UCG GGC UUC GGU CCG GUU C | |
| *btuB*_SHAPE | GGC CGG UCC UGU GAG GUU AAU AGG GAA UCC AGU GCG AAU CUG GAG CUG ACG CGC AGC UGU UAG GAA AGG UGC CAU GAU GUC GUU AUG CGG ACA CUC GCC AUU CGG UGG GAA GUC AUC AUG UCU UAG UAU CUU AGA UAC CCC UCC AAG CCA GGA GAC CUG CCG GCC AAC GUC GCA UCU GGU UCU CAU CAU CGC GUA AUA UUG AUG AAA CCU GCG GCA UCG AUC CGG UUC GCC GGA UCC AAA UCG GGC UUC GGU CCG GUU C | |
| SHAPE reverse transcription (RT) primer | GAA CCG GAC CGA AGC CCG | |
| inner RT primer for *btuB*_SHAPE | GAA CCG GAC CGA AGC CCG ATT GCC GGC AGG TCT TCG GGC TTG GA | |
| *env8*_A20U | GGC CUA AAA GCG UAG UGG GUA AGU GAC GUG AAA UUC GUC CAG AUU ACU UGA UAC | |

| | |
|---|---|
| | GGU UAU ACU CCG AAU GCC ACC UAG GCC AUA CAA CGA GCA AGG AGA CUC |
| *env8*_A20C | GGC CUA AAA GCG UAG UGG GCA AGU GAC GUG AAA UUC GUC CAG AUU ACU UGA UAC GGU UAU ACU CCG AAU GCC ACC UAG GCC AUA CAA CGA GCA AGG AGA CUC |
| *env8*_A20G | GGC CUA AAA GCG UAG UGG GGA AGU GAC GUG AAA UUC GUC CAG AUU ACU UGA UAC GGU UAU ACU CCG AAU GCC ACC UAG GCC AUA CAA CGA GCA AGG AGA CUC |
| *env8*_G19U | GGC CUA AAA GCG UAG UGG UAA AGU GAC GUG AAA UUC GUC CAG AUU ACU UGA UAC GGU UAU ACU CCG AAU GCC ACC UAG GCC AUA CAA CGA GCA AGG AGA CUC |
| *env8*_A21U | GGC CUA AAA GCG UAG UGG GAU AGU GAC GUG AAA UUC GUC CAG AUU ACU UGA UAC GGU UAU ACU CCG AAU GCC ACC UAG GCC AUA CAA CGA GCA AGG AGA CUC |
| *env8*_J6/3 | GGC CUA AAA GCG UAG UGG GAA AGU GAC GUG AAA UUC GUC CAG AUU ACU UGA UAC GGU UAU ACU CCG AGU GCC ACC UAG GCC AUA CAA CGA GCA AGG AGA CUC |
| *env8*_J6/3+A20G | GGC CUA AAA GCG UAG UGG GGA AGU GAC GUG AAA UUC GUC CAG AUU ACU UGA UAC GGU UAU ACU CCG AGU GCC ACC UAG GCC AUA CAA CGA GCA AGG AGA CUC |
| *env8*_J1/3 | GGC CUA AAU CAU AGU GGG AAA GUG ACG UGA AAU UCG UCC AGA UUA CUU GAU ACG GUU AUA CUC CGA AUG CCA CCU AGG CCA UAC AAC GAG CAA GGA GAC UC |
| *env8*_J1/3+J6/3+A20G | GGC CUA AAU CAU AGU GGG GAA GUG ACG UGA AAU UCG UCC AGA UUA CUU GAU ACG GUU AUA CUC CGA GUG CCA CCU AGG CCA UAC AAC GAG CAA GGA GAC UC |

**Table S2:** Degree of protection values as plotted in Figures 2, 3, S2, S3, and S4.

| Figure | RNA | Region | Cbl | Rep 1 | Rep 2 | Rep 3 | Rep 4 | Rep 5 | Rep 6 | Rep 7 | Rep 8 | Rep 9 | Rep 10 |
|---|---|---|---|---|---|---|---|---|---|---|---|---|---|
| 2A | *env8* | L5 | CNCbl | 7.9E-01 | 8.7E-01 | 8.6E-01 | 8.9E-01 | 6.3E-01 | 6.2E-01 | 6.8E-01 | | | |
| | | | *i* | 7.5E-01 | 8.0E-01 | 7.3E-01 | 8.3E-01 | 5.4E-01 | 5.0E-01 | 5.1E-01 | | | |
| | | | *ii* | 6.9E-01 | 7.0E-01 | 5.9E-01 | 7.9E-01 | 5.7E-01 | 5.2E-01 | 5.7E-01 | | | |
| | | | *iii* | 6.7E-01 | 6.8E-01 | 6.0E-01 | 7.2E-01 | 5.1E-01 | 4.8E-01 | 4.8E-01 | | | |
| | | | *iv* | 3.2E-01 | 6.4E-02 | -8.3E-02 | 8.6E-02 | -7.5E-02 | 1.5E-02 | -1.7E-02 | | | |
| | | | *v* | 5.7E-01 | 4.3E-01 | 4.0E-02 | 4.5E-01 | 3.2E-01 | 3.1E-01 | 1.5E-01 | | | |
| | | | *vi* | 4.8E-01 | 3.8E-01 | 2.3E-02 | 3.0E-01 | 3.1E-01 | 1.6E-01 | | | | |
| | | | *vii* | 4.7E-01 | 9.3E-02 | 3.3E-01 | 1.3E-01 | 1.1E-01 | 2.0E-01 | | | | |
| 2B | *env8* | L5 | CNCbl | 7.9E-01 | 8.7E-01 | 8.6E-01 | 8.9E-01 | 7.0E-01 | 7.1E-01 | 6.2E-01 | | | |
| | | | *viii* | 4.5E-01 | 6.5E-01 | 6.1E-01 | 7.5E-01 | 5.0E-01 | 5.0E-01 | 4.7E-01 | | | |
| | | | *ix* | 7.8E-01 | 8.1E-01 | 7.9E-01 | 8.3E-01 | 5.9E-01 | 6.0E-01 | 5.4E-01 | | | |
| | | | *x* | 7.3E-01 | 8.1E-01 | 7.7E-01 | 8.4E-01 | 5.3E-01 | 5.6E-01 | 5.1E-01 | | | |
| | | | *xi* | 4.3E-01 | 5.1E-01 | 1.6E-01 | 5.5E-01 | 2.1E-01 | 2.1E-01 | 3.6E-01 | | | |
| | | A20 | CNCbl | 7.4E-01 | 7.7E-01 | 8.1E-01 | 8.0E-01 | 5.7E-01 | 5.4E-01 | 5.6E-01 | 5.0E-01 | 4.7E-01 | 5.0E-01 |
| | | | *viii* | -7.6E-01 | -1.3E+00 | -1.4E+00 | -8.1E-01 | -1.1E+00 | -1.4E+00 | -8.0E-01 | | | |
| | | | *ix* | -1.1E+00 | -1.7E+00 | -2.3E+00 | -1.4E+00 | -1.8E+00 | -1.9E+00 | -1.1E+00 | | | |
| | | | *x* | 5.0E-01 | 4.5E-01 | 3.7E-01 | 5.6E-01 | -1.4E-01 | -1.3E-01 | 1.2E-01 | | | |
| | | | *xi* | 3.9E-01 | 3.0E-01 | 2.8E-02 | 4.0E-01 | -2.8E-01 | -2.0E-01 | -1.2E-02 | | | |
| 3A | *env50* | J6/3 | CNCbl | 1.4E-02 | 2.0E-02 | 2.4E-02 | | | | | | | |
| | | | AdoCbl | 1.1E-01 | 1.6E-01 | 2.1E-01 | | | | | | | |
| | | | *viii* | 2.1E-02 | 3.0E-02 | 3.0E-02 | | | | | | | |
| | | | *ix* | 1.1E-02 | 2.1E-02 | 2.2E-02 | | | | | | | |
| | | | *x* | 1.7E-02 | 2.1E-02 | 2.1E-02 | | | | | | | |
| | | | *xi* | 4.2E-03 | 1.4E-02 | 1.8E-02 | | | | | | | |
| | | J1/13 | CNCbl | -3.2E-02 | -6.5E-03 | -5.7E-02 | | | | | | | |
| | | | AdoCbl | -1.5E-01 | -1.3E-01 | -2.8E-01 | | | | | | | |
| | | | *viii* | 1.8E-02 | 1.2E-02 | -2.0E-02 | | | | | | | |
| | | | *ix* | 8.5E-03 | 6.7E-02 | 4.1E-03 | | | | | | | |

| Figure | RNA | Region | Cbl | Rep 1 | Rep 2 | Rep 3 | Rep 4 | Rep 5 | Rep 6 | Rep 7 | Rep 8 | Rep 9 | Rep 10 |
|---|---|---|---|---|---|---|---|---|---|---|---|---|---|
| | | | *x* | 2.4E-02 | 7.1E-02 | -2.2E-03 | | | | | | | |
| | | | *xi* | 1.8E-02 | 5.9E-02 | 1.1E-02 | | | | | | | |
| 3B | *btuB* | L5 | AdoCbl | 5.9E-01 | 6.6E-01 | 6.0E-01 | | | | | | | |
| | | | *viii* | 8.9E-02 | 1.9E-02 | 1.8E-01 | | | | | | | |
| | | | *ix* | 3.0E-02 | -9.3E-02 | 9.9E-04 | | | | | | | |
| | | | *x* | 1.1E-01 | 2.0E-01 | 2.0E-01 | | | | | | | |
| | | | *xi* | -1.6E-02 | 4.2E-02 | 1.7E-02 | | | | | | | |
| | | | CNCbl | 1.9E-01 | 4.1E-01 | -1.6E-01 | | | | | | | |
| | | J11/10 | AdoCbl | 5.6E-01 | 5.4E-01 | 5.5E-01 | | | | | | | |
| | | | *viii* | -1.5E-01 | -1.8E-02 | -1.3E-01 | | | | | | | |
| | | | *ix* | -7.1E-02 | -6.3E-03 | -6.4E-02 | | | | | | | |
| | | | *x* | -1.1E-01 | 7.8E-02 | -9.1E-03 | | | | | | | |
| | | | *xi* | -1.5E-01 | 3.9E-02 | -1.7E-01 | | | | | | | |
| | | | CNCbl | -8.0E-02 | 6.1E-02 | -2.4E-01 | | | | | | | |
| S2C | *env8* | J6/3 | CNCbl | 7.2E-01 | 7.4E-01 | 7.9E-01 | 7.6E-01 | 4.3E-01 | 4.4E-01 | 4.5E-01 | 4.6E-01 | 4.8E-01 | 4.1E-01 |
| | | | *i* | 6.1E-01 | 6.3E-01 | 5.5E-01 | 7.3E-01 | 2.6E-01 | 2.2E-01 | 1.8E-01 | | | |
| | | | *ii* | 8.5E-01 | 6.2E-01 | 5.4E-01 | 7.2E-01 | 4.2E-01 | 3.6E-01 | 3.4E-01 | | | |
| | | | *iii* | 5.6E-01 | 6.0E-01 | 5.1E-01 | 6.5E-01 | 3.8E-01 | 3.5E-01 | 3.2E-01 | | | |
| | | | *iv* | -4.3E-01 | 5.7E-02 | 4.2E-01 | 2.1E-01 | -1.3E-02 | 2.9E-02 | -8.5E-02 | | | |
| | | | *v* | 4.1E-01 | 3.1E-01 | 5.2E-01 | 4.2E-01 | 2.0E-01 | 1.9E-01 | 4.7E-02 | | | |
| | | | *vi* | 3.0E-01 | 2.0E-01 | 4.0E-01 | 1.9E-01 | 1.8E-01 | 4.6E-03 | | | | |
| | | | *vii* | 2.5E-01 | 6.3E-02 | -2.7E-01 | 1.3E-01 | 1.0E-01 | -5.3E-02 | | | | |
| | | | *viii* | 3.4E-01 | 6.1E-01 | 6.6E-01 | 7.0E-01 | 3.1E-01 | 3.0E-01 | 3.2E-01 | | | |
| | | | *ix* | 7.4E-01 | 8.1E-01 | 8.1E-01 | 8.0E-01 | 4.3E-01 | 4.6E-01 | 4.1E-01 | | | |
| | | | *x* | 6.8E-01 | 7.6E-01 | 7.3E-01 | 7.8E-01 | 3.4E-01 | 3.9E-01 | 3.3E-01 | | | |
| | | | *xi* | 3.5E-01 | 4.6E-01 | 5.3E-01 | 5.4E-01 | 1.1E-01 | 1.7E-01 | 2.3E-01 | | | |
| | | J1/13 | CNCbl | -6.5E-01 | -3.4E-01 | -4.8E-02 | -1.9E-01 | -2.2E-01 | -1.0E-01 | | | | |
| | | | *i* | -5.7E-01 | -2.8E-01 | -1.4E-01 | -1.8E-01 | -2.6E-01 | -5.7E-01 | | | | |
| | | | *ii* | -9.2E-02 | -2.7E-01 | -2.0E-01 | -1.5E-01 | -2.9E-01 | -4.4E-01 | | | | |

| Figure | RNA | Region | Cbl | Rep 1 | Rep 2 | Rep 3 | Rep 4 | Rep 5 | Rep 6 | Rep 7 | Rep 8 | Rep 9 | Rep 10 |
|---|---|---|---|---|---|---|---|---|---|---|---|---|---|
| | | | *iii* | -2.9E-01 | -2.3E-01 | -1.4E-01 | -2.7E-01 | -3.0E-01 | -4.8E-01 | | | | |
| | | | *iv* | 4.1E-02 | 7.8E-02 | 2.8E-01 | -1.4E-02 | -1.8E-02 | -7.0E-02 | | | | |
| | | | *v* | -2.6E-01 | -1.6E-01 | -8.9E-02 | -3.0E-01 | -3.2E-01 | -4.7E-01 | | | | |
| | | | *vi* | -4.7E-02 | -2.2E-01 | -4.5E-02 | -5.3E-02 | -3.4E-01 | | | | | |
| | | | *vii* | 1.5E-01 | -5.2E-02 | 5.5E-02 | 1.9E-02 | -2.5E-01 | | | | | |
| | | | *viii* | -2.9E-01 | -2.8E-01 | -1.5E-02 | -1.9E-01 | -2.7E-01 | -1.2E-01 | | | | |
| | | | *ix* | -3.3E-01 | -3.2E-01 | -1.6E-01 | -4.1E-01 | -5.4E-01 | -2.4E-01 | | | | |
| | | | *x* | -4.0E-01 | -1.9E-01 | -3.9E-02 | -3.0E-01 | -3.6E-01 | -3.7E-01 | | | | |
| | | | *xi* | -8.5E-02 | -2.0E-01 | 3.7E-03 | -1.1E-01 | -1.5E-01 | -1.8E-01 | | | | |
| S3C | *env50* | J6/3 | CNCbl | 8.7E-02 | 9.5E-02 | 2.1E-01 | | | | | | | |
| | | | AdoCbl | 4.3E-01 | 4.6E-01 | 5.8E-01 | | | | | | | |
| | | | *ii* | 5.9E-02 | 1.8E-02 | 1.2E-01 | | | | | | | |
| | | | *i* | 9.6E-02 | 1.9E-01 | -1.2E-02 | | | | | | | |
| | | | *iii* | 8.8E-02 | 9.7E-02 | -1.2E-02 | | | | | | | |
| | | | *iv* | 3.9E-02 | 3.9E-02 | 6.4E-02 | | | | | | | |
| | | | *v* | 6.5E-02 | 6.1E-02 | 9.8E-03 | | | | | | | |
| | | | *vi* | 1.1E-01 | 3.3E-02 | -6.9E-02 | | | | | | | |
| | | | *vii* | 1.0E-01 | 1.4E-01 | 9.5E-03 | | | | | | | |
| | | J1/13 | CNCbl | -5.5E-02 | -1.2E-01 | -1.3E-01 | | | | | | | |
| | | | AdoCbl | 2.9E-02 | 2.9E-02 | 2.9E-02 | | | | | | | |
| | | | *ii* | 2.9E-02 | 2.9E-02 | 2.9E-02 | | | | | | | |
| | | | *i* | -5.7E-02 | -1.6E-01 | -5.1E-02 | | | | | | | |
| | | | *iii* | 3.7E-03 | -5.5E-02 | 9.1E-03 | | | | | | | |
| | | | *iv* | -2.0E-02 | -1.5E-02 | -2.8E-02 | | | | | | | |
| | | | *v* | -2.0E-02 | -1.3E-01 | 2.9E-02 | | | | | | | |
| | | | *vi* | -4.1E-02 | -1.3E-01 | -7.8E-02 | | | | | | | |
| | | | *vii* | 6.2E-03 | -9.5E-02 | 3.2E-02 | | | | | | | |
| S4C | *btuB* | L5 | AdoCbl | 6.0E-01 | 5.5E-01 | 4.0E-01 | | | | | | | |
| | | | *i* | -4.0E-02 | -8.4E-02 | -2.2E-02 | | | | | | | |

| **Figure** | **RNA** | **Region** | **Cbl** | **Rep 1** | **Rep 2** | **Rep 3** | **Rep 4** | **Rep 5** | **Rep 6** | **Rep 7** | **Rep 8** | **Rep 9** | **Rep 10** |
|---|---|---|---|---|---|---|---|---|---|---|---|---|---|
| | | | *ii* | 8.7E-03 | 1.0E-01 | 5.7E-02 | | | | | | | |
| | | | *iii* | -1.4E-01 | -7.9E-02 | -2.7E-01 | | | | | | | |
| | | | *iv* | 3.4E-02 | -2.1E-01 | -1.0E-01 | | | | | | | |
| | | | *v* | 6.2E-02 | -1.7E-01 | -2.7E-02 | | | | | | | |
| | | | *vi* | -6.5E-02 | -1.8E-01 | -3.2E-02 | | | | | | | |
| | | | *vii* | 5.2E-02 | -1.1E-01 | 1.9E-01 | | | | | | | |
| | | | CNCbl | 1.3E-01 | -1.8E-01 | 3.3E-02 | | | | | | | |
| | | J11/10 | AdoCbl | 5.7E-01 | 4.3E-01 | 4.7E-01 | | | | | | | |
| | | | *i* | -8.7E-02 | -1.2E-01 | 8.9E-03 | | | | | | | |
| | | | *ii* | -1.7E-02 | -4.8E-02 | 1.2E-02 | | | | | | | |
| | | | *iii* | -2.4E-02 | -1.3E-01 | 6.0E-02 | | | | | | | |
| | | | *iv* | -8.0E-03 | -1.3E-01 | 1.1E-02 | | | | | | | |
| | | | *v* | -2.6E-02 | -1.9E-01 | 3.0E-02 | | | | | | | |
| | | | *vi* | -1.1E-02 | -1.1E-01 | 5.3E-02 | | | | | | | |
| | | | *vii* | -2.1E-03 | -9.6E-02 | 7.0E-02 | | | | | | | |
| | | | CNCbl | 1.9E-02 | -1.1E-01 | 2.0E-02 | | | | | | | |

### Supplemental Methods

### Synthesis and characterization of the cobalamin derivatives and probes

### General Information

Commercially available reagents and solvents were used as received. ATTO propargylamide was obtained from ATTO-TEC. As supplied ATTO 590 consists of a mixture of two isomers with practically identical absorption and fluorescence (para and meta isomer). For simplicity only one isomer is presented on the schemes. The scale of the reaction with ATTO dye did not provide sufficient amount of the product for NMR analyses, thus the HPLC and MS analyses were performed to characterize and confirm the purity of the conjugate. $^{1}$H and $^{13}$C NMR spectra were recorded at rt on a Bruker 400 MHz spectrometer with the residual solvent peak used as an internal standard. Data are reported as follows: chemical shift, peak multiplicity (s = singlet, d = doublet, t = triplet, q = quartet, m = multiplet), coupling constants (Hz), and number of protons. High-resolution ESI mass spectra were recorded on Waters Synapt G2 HDMS qTOF. All reactions and product purities were monitored using RP-HPLC techniques. Preparative chromatography was performed using LiChroprep RP-18 (40–63 mm) with HPLC grade water and MeCN as eluents. HPLC measurement conditions: column, Kromasil 100-5-C18, 250 mm, 4.6 mm; detection, UV/Vis; pressure, 20 MPa; temperature, 22ºC. Abbreviations: AcOEt – ethyl acetate; Azido-PEG5-amine – 17-Azido-3,6,9,12,15-pentaoxaheptadecan-1-amine; CDT – 1,1′-Carbonyl-di-(1,2,4-triazole); $Et_2O$ – diethyl ether; MeCN – acetonitrile; MeOH – methanol; NMP – *N*-Methyl-2-pyrrolidone; RP HPLC – Reverse-phase high-performance liquid chromatography; TBTA – Tris[(1-benzyl-1H-1,2,3-triazol-4 yl)methyl]amine; TEA – Triethylamine; TFA – trifluoroacetic acid.

### Synthesis of cobalamin acetylides (*viii-xi*)

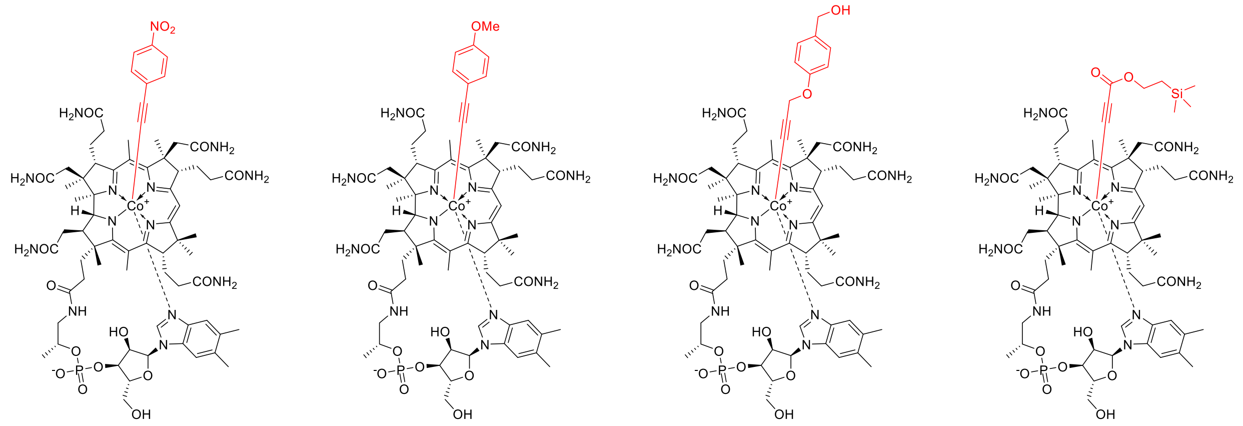

All of the above derivatives were synthesized according to described procedures.[2-3] All spectra matched those reported in the literature.

**Synthesis of cobalamin acetylide *viii* azide**

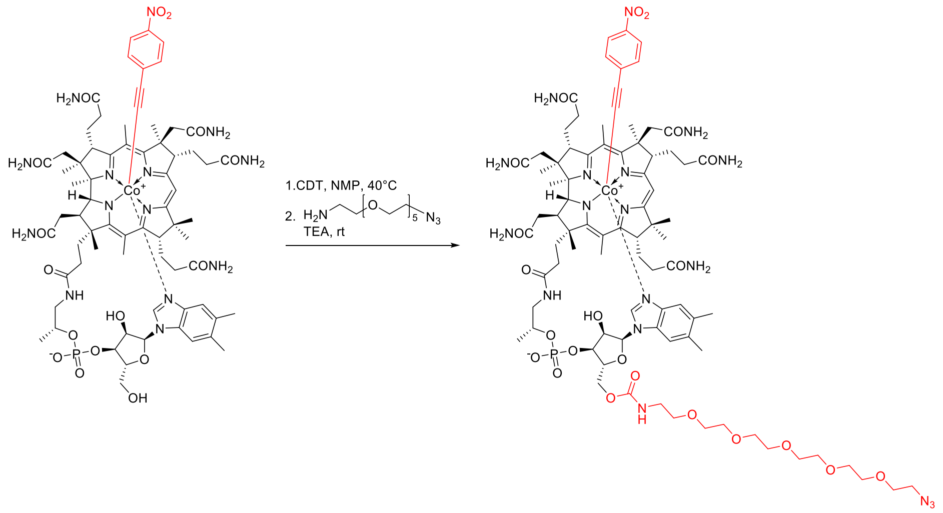

Acetylide *viii* (30 mg, 0.02 mmol, 1 equiv.) was dissolved in dry *N*-Methyl-2-pyrrolidone (NMP, 1.5 mL) at 40 °C under an argon atmosphere. To a stirring solution under argon solid CDT (6.6 mg, 0.04 mmol, 2 equiv.) was added. When full consumption of the substrate (monitored by the RP HPLC) was observed (approx. 1.5 h), heating bath was removed and azido-PEG5-amine (25 mg, 0.08 mmol, 4 equiv.) was added in one portion along with TEA (4.0 mg, 0.04 mmol, 2 equiv.). The resulting solution was stirred overnight. Subsequently, the reaction mixture was poured into AcOEt (10 mL), and centrifuged. The solid residue was redissolved in MeOH and precipitated with $Et_2O$ (10 mL). After drying, the remaining solid was dissolved in water and purified by RP column chromatography (60 mL) with a mixture of MeCN and $H_2O$ as eluents (gradually from 10% to 20% v/v). The desired compound was obtained as a red powder; yield: 75%. $^1$H NMR (400 MHz, $CD_3OD$) δ 7.97 (d, *J* = 9.0 Hz, 2H), 7.24 (s, 1H), 7.23 (s, 1H), 7.00 (d, *J* = 9.0 Hz, 2H), 6.62 (s, 1H), 6.23 (d, *J* = 2.7 Hz, 1H), 6.00 (s, 1H), 4.67 (dd, *J* = 11.9, 1.9 Hz, 1H), 4.64 – 4.56 (m, 1H), 4.42 – 4.31 (m, 2H), 4.30 – 4.21 (m, 2H), 4.17 (dd, *J* = 11.9, 2.1 Hz, 1H), 3.70 – 3.59 (m, 11H), 3.64 (s, 3H), 3.629 (s, 3H), 3.626 (s, 3H), 3.56 – 3.51 (m, 2H), 3.38 – 3.34 (m, 2H), 3.30 – 3.23 (m, 2H), 3.00 – 2.90 (m, 1H), 2.88 – 2.80 (m, 1H), 2.64 – 1.83 (m, 22H), 2.60 (s, 3H), 2.57 (s, 3H), 2.29 (s, 3H), 2.29 (s, 3H), 1.85 (s, 3H), 1.79 – 1.70 (m, 1H), 1.48 (s, 3H), 1.36 (s, 3H), 1.31 (s, 3H), 1.34 – 1.27 (m, 1H),1.24 (d, *J* = 6.3 Hz, 3H), 1.12 (s, 3H), 0.53 (s, 3H). $^{13}$C NMR (100 MHz, $CD_3OD$) δ 180.0, 178.4, 177.6, 177.4, 176.8, 176.2, 175.9, 175.6, 174.8, 174.24, 174.17, 166.5, 166.1, 158.7, 146.6, 143.6, 138.7, 135.1, 134.6, 133.4, 132.5, 131.5, 124.5, 118.5, 112.1, 108.0, 104.9, 102.6, 95.4, 88.1, 86.2, 81.2, 75.8, 75.3, 73.4, 71.6, 71.5, 71.3, 71.1, 71.0, 70.6, 66.9, 64.3, 59.9, 57.0, 56.3, 55.1, 52.2, 51.8, 46.5, 43.7, 43.5, 41.8, 40.0, 36.4, 35.2, 33.1, 32.7, 32.5, 32.0, 29.5, 27.5, 27.4, 20.9, 20.8, 20.42, 20.37, 20.2, 20.1, 19.9, 17.5, 17.2, 16.4, 16.2, 15.4. HRMS

(ESI) m/z [M + H]$^+$ calculated for $C_{83}H_{117}CoN_{18}O_{22}P^+$, 1807.7655; found, 1807.7656. $t_R$ (RP-HPLC, from 10% MeCN/$H_2O$ + 0.05% TFA to 70% MeCN/$H_2O$ + 0.05% TFA in 16 min): 9.76 min.

$^1$H NMR spectrum (400 MHz, $CD_3OD$)

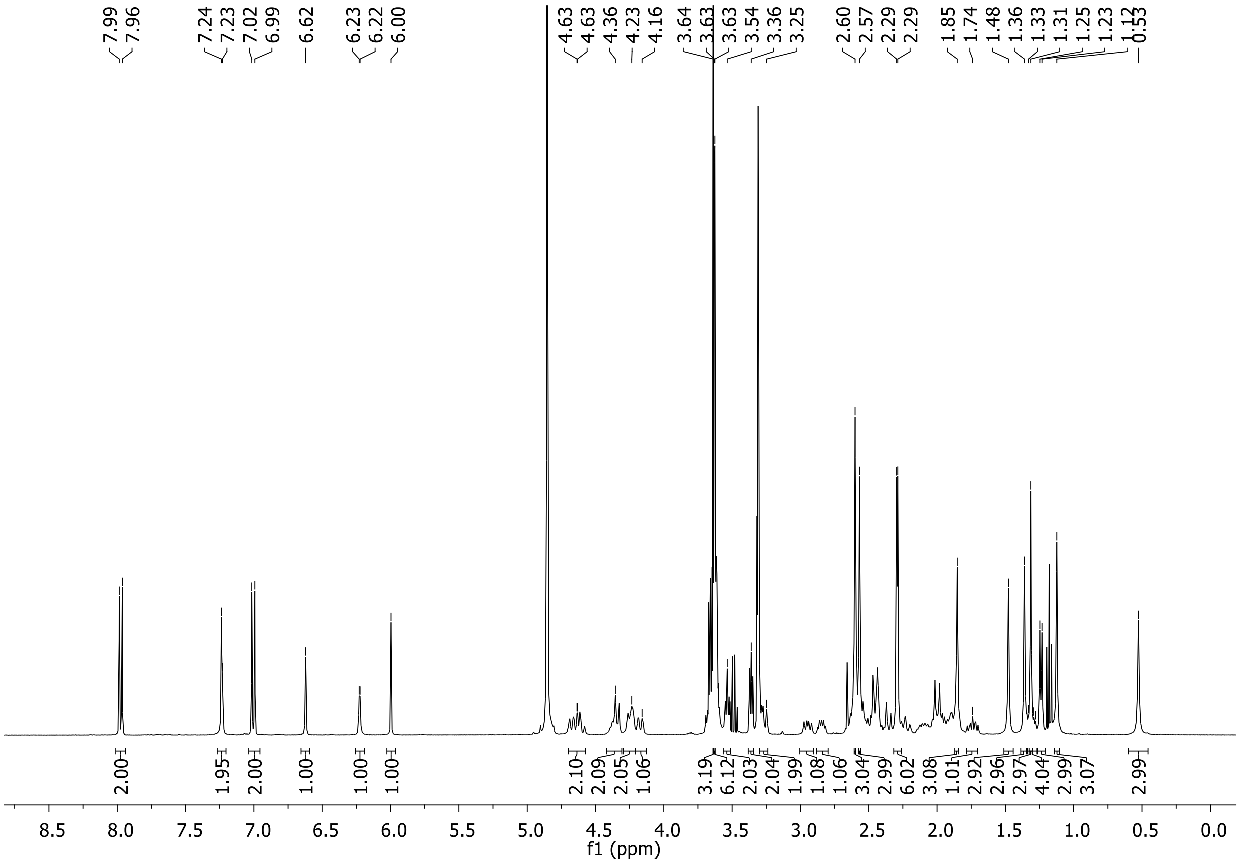

$^{13}$C NMR spectrum (100 MHz, $CD_3OD$)

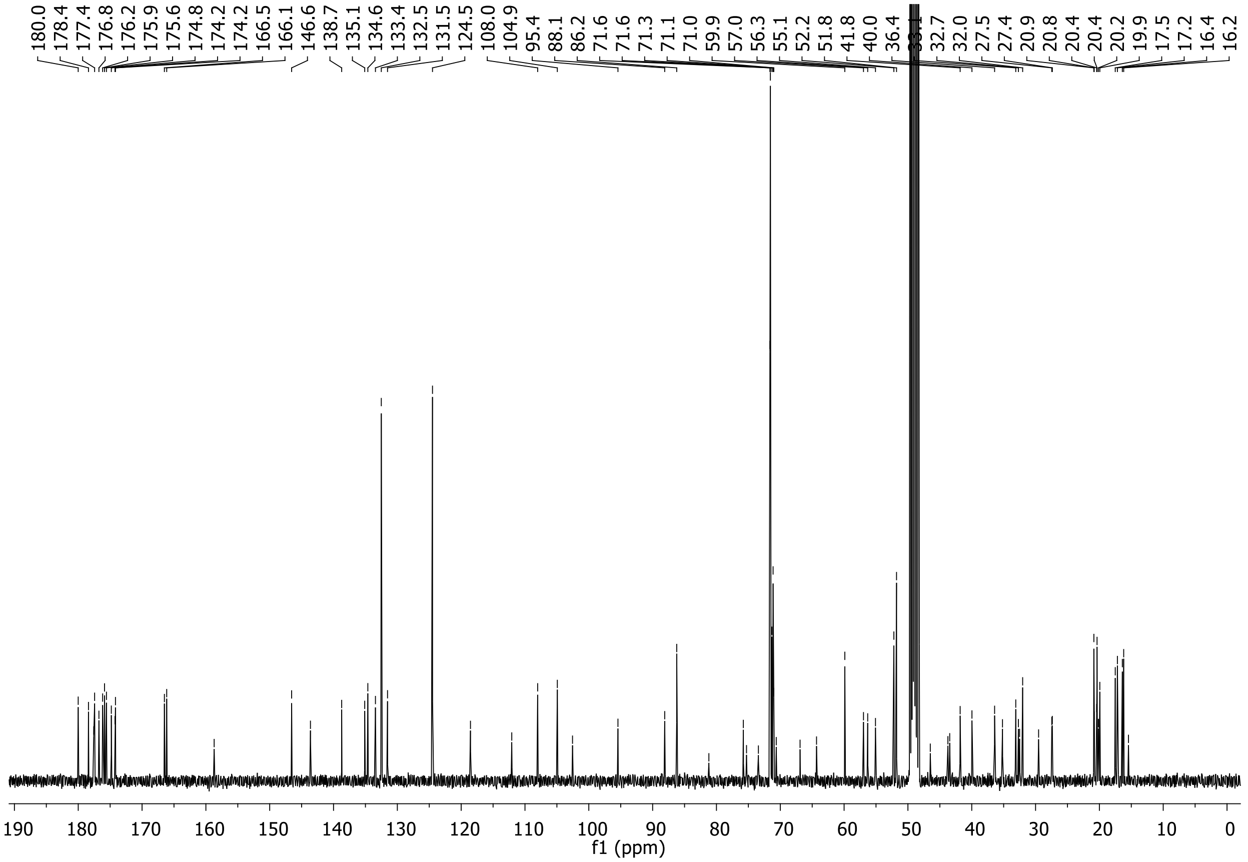

### HPLC chromatogram (λ= 361 nm)

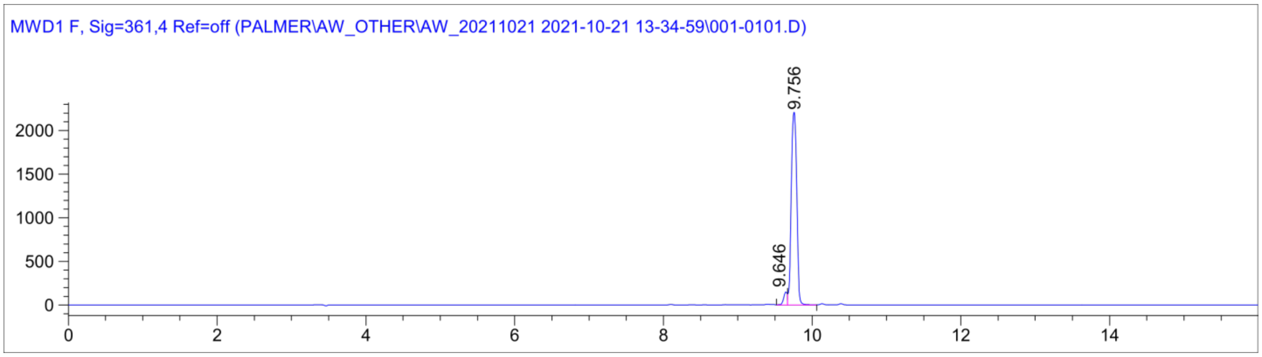

### ESI MS spectrum

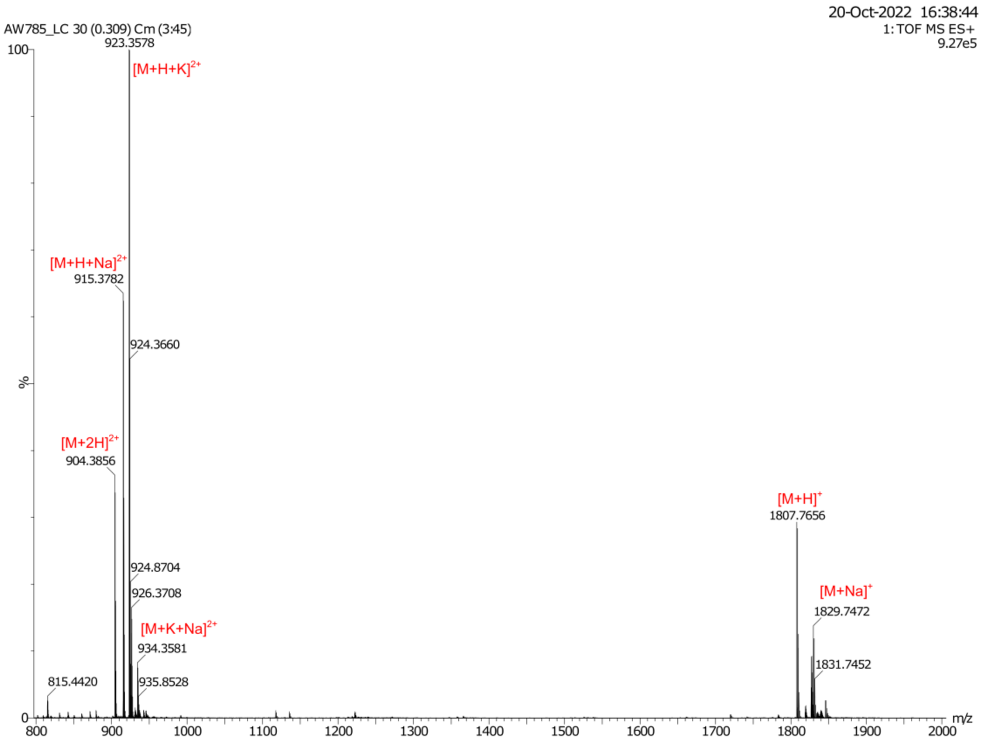

**Synthesis of cobalamin acetylide *viii*-dye conjugate, the PhNO2-Cbl-5xPEG-ATTO590 probe**

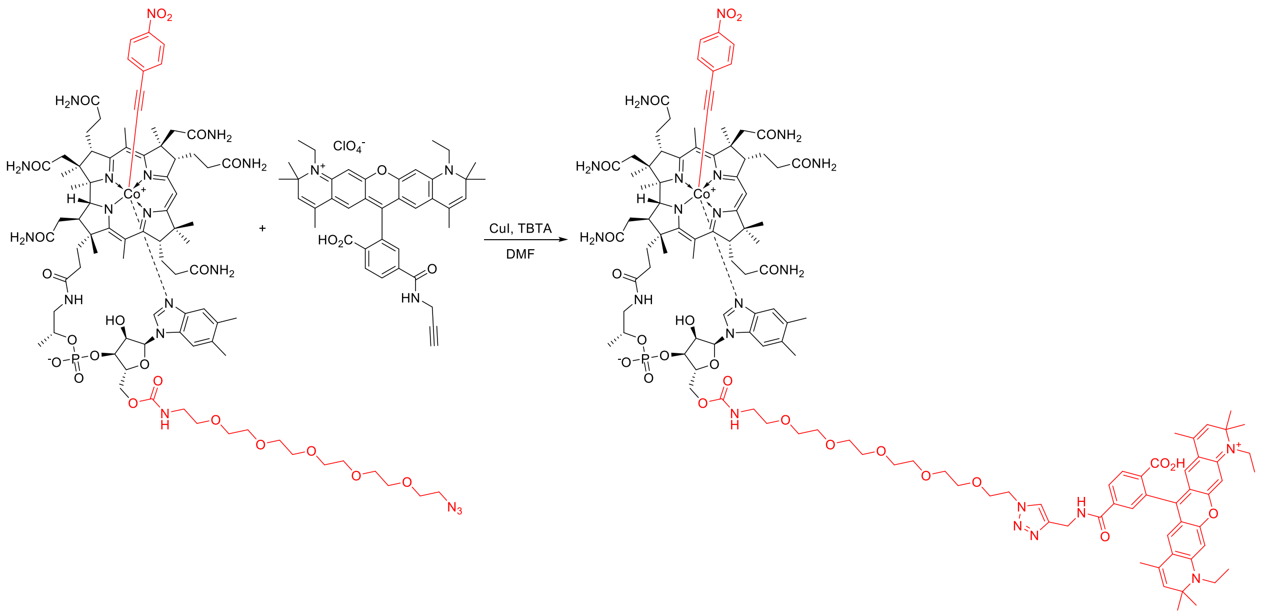

Preparation of a catalyst solution: CuI (1 mg, 5 μmol) and TBTA (5 mg, 10 μmol) were dissolved in DMF (1 mL) and stirred for 20 min. Acetylide *viii* azide (2.5 mg, 1.38 μmol, 2 equiv.) and ATTO 590 alkyne (0.5 mg, 0.69 μmol, 1 equiv.) were dissolved in DMF (0.3 mL). Subsequently, freshly prepared catalyst solution (300 μL) was added and the reaction mixture was stirred overnight (full consumption of the dye was confirmed via RP HPLC). The reaction mixture was poured into AcOEt (10 mL) and the precipitate was centrifuged and dried. The crude solid was subsequently redissolved in MeOH (1 mL), precipitated with $Et_2O$ (10 mL) and centrifuged. The dried solid was then dissolved in $H_2O$, loaded onto RP column (10 mL) and purified gradually with MeCN/$H_2O$ from 15 to 35% v/v yielding product as a violet solid. HRMS (ESI) m/z $[M + H]^{2+}$ calcd for $C_{123}H_{159}CoN_{21}O_{26}P^{2+}$, 1218.0412; found, 1218.0414. $t_R$ (RP-HPLC, from 10% MeCN/$H_2O$ + 0.05% TFA to 70% MeCN/$H_2O$ + 0.05% TFA in 16 min): 11.17 min.

HPLC chromatogram ($\lambda$= 590 nm)

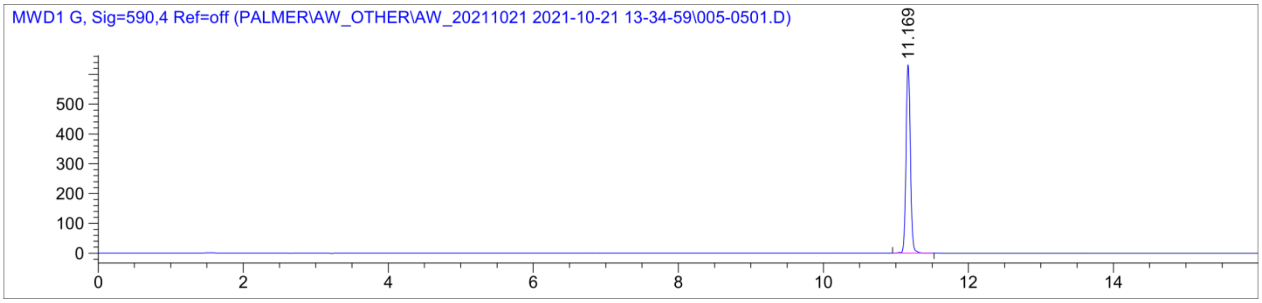

ESI MS spectrum

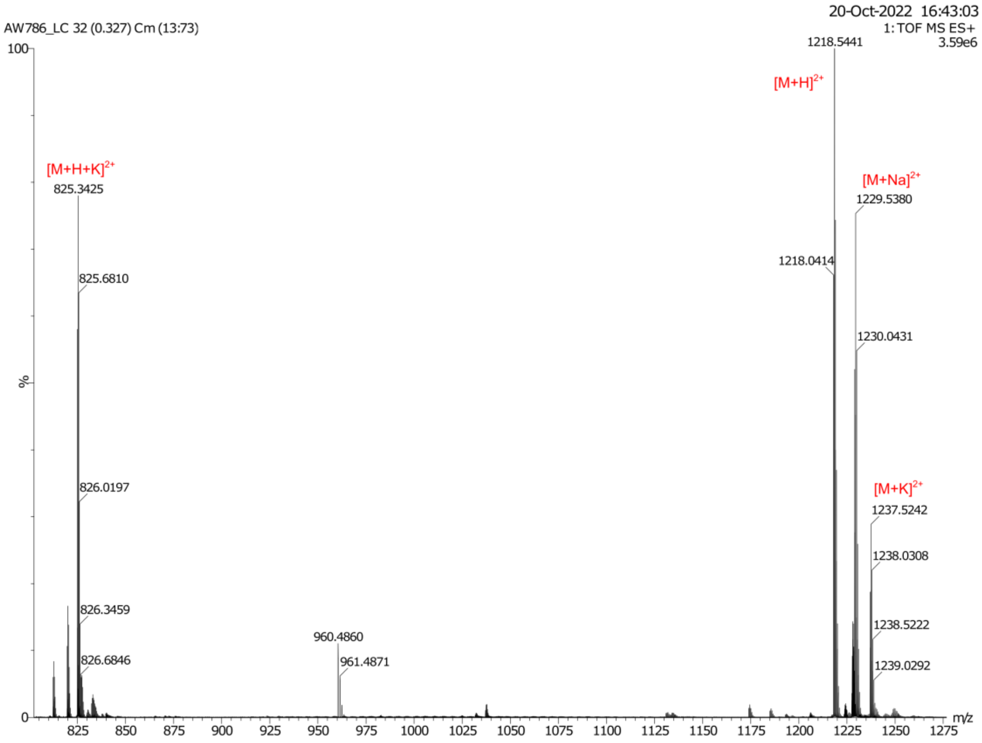

**Synthesis of *meso*-modified cobalamins (*iii-vii*)**

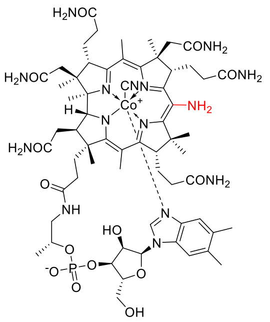

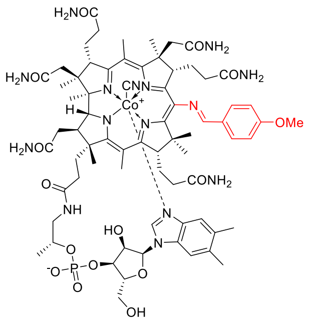

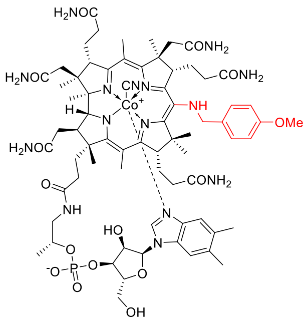

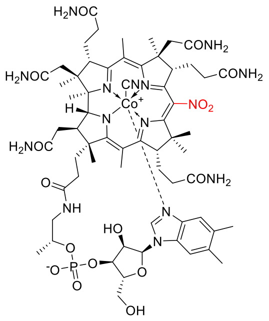

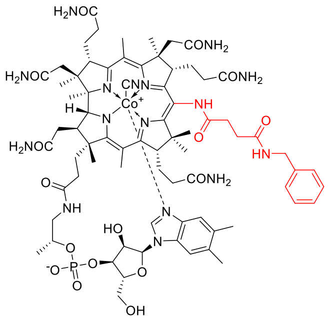

All of the above derivatives were synthesized according to described procedures.[4] All spectra matched those reported in the literature.

#### Synthesis of cobalamin lactones (*i-ii*)

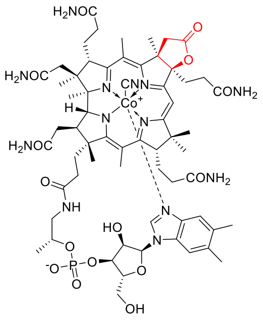

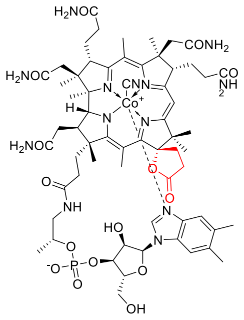

Cobalamin *c*-lactone[5] and cobalamin *e*-lactone[4] were synthesized according to described procedures. All spectra matched those reported in the literature.

#### Selective 2'-Hydroxyl Acylation Analyzed by Primer Extension (SHAPE)

Changes to described procedure are as follows: *E. coli btuB* Cbl riboswitch RNA was refolded by incubating at 65 °C for 3 minutes, room temperature for 5 minutes, and on ice for a minimum of five minutes. All other RNAs were refolded by heating at 90 °C for 3 minutes and placing on ice for a minimum of ten minutes. RNA was added to folding buffer and ligand and incubated at room temperature for 10 minutes before adding NMIA or neat DMSO for control reactions. The final 10 μL probing reactions had concentrations of 100 nM RNA, 100 mM Na-HEPES pH 8.0, 100 mM NaCl, 6 mM MgCl2, 30 μM ligand unless otherwise noted, and 13 mM NMIA. Probing reactions were incubated at 37 °C for 41 minutes (5 half-lives). Following probing, reactions were moved straight into primer extension as previously described.[6] Products were run on a 12% denaturing polyacrylamide gel and visualized using a Typhoon FLA 9500 PhosphoroImager (GE).

Degree of protection for indicated areas of each RNA were calculated using the SAFA software package.[7] The software was used to align gel images, and band positions were manually assigned using the sequencing lanes as a guide before quantification of band intensities. Invariant residues of replicate gels were determined by the software, and the band intensity values for each gel were normalized to a set of invariant residues common to all replicates. The intensities for every band in a region of interest were added, and region intensities for each ligand condition were normalized to those of the NMIA condition. This resulted in the degree of protection values plotted in the figures. Degree of protection values for each individual replicate can be found in **Table S2**.

**Supplemental References.**
